# Modeling of blood flow dynamics in the rat somatosensory cortex

**DOI:** 10.1101/2024.11.14.623572

**Authors:** Stéphanie Battini, Nicola Cantarutti, Christos Kotsalos, Yann Roussel, Alessandro Cattabiani, Alexis Arnaudon, Cyrille Favreau, Stefano Antonel, Henry Markram, Daniel Keller

## Abstract

The cerebral microvasculature forms a dense network of interconnected blood vessels where flow is modulated partly by astrocytes. Increased neuronal activity stimulates astrocytes to release vasoactive substances at the endfeet, altering the diameters of connected vessels. Our study simulated the coupling between blood flow variations and vessel diameter changes driven by astrocytic activity in the rat somatosensory cortex. We developed a framework with three key components: coupling between vasculature and synthesized astrocytic morphologies, a fluid dynamics model to compute flow in each vascular segment, and a stochastic process replicating the effect of astrocytic endfeet on vessel radii. The model was validated against experimental flow values from literature across cortical depths. We found that local vasodilation from astrocyte activity increased blood flow, especially in capillaries, exhibiting a layer-specific response in deeper cortical layers. Additionally, the highest blood flow variability occurred in capillaries, emphasizing their role in cerebral perfusion regulation. We discovered that astrocytic activity impacts blood flow dynamics in a localized, clustered manner, with most vascular segments influenced by two to three neighboring endfeet. These insights enhance our understanding of neurovascular coupling and guide future research on blood flow-related diseases.

## 1. Introduction

The interrelationships among astrocytes, neurons, and blood vessels are particularly complex, giving rise to a system of multidirectional communication, the neuro-glia-vascular (NGV) ensemble [1]. Astrocytes play an important role in it and in energy metabolism [2]. Of particular note are the specialized astrocytic processes known as *endfeet*, which establish direct contact with arterioles, venules, and capillaries, securely anchoring them to the vascular surface [3–5]. Several investigations relying on chemical fixation for tissue preservation reported that astrocytic endfeet cover at least 60% of the total surface area of blood vessels [3,5–7]. In accordance to that, we build upon the recent *in silico* NGV circuit developed by Zisis et al. [8] that exhibits a vascular coverage of approximately 60%.

Astrocytes regulate some aspects of blood flow and the exchange of molecules to and from the brain. This regulation is partially coordinated by neuronal activity monitored by astrocytes and mediated via their endfeet. Release of substances from interneurons, such as nitric oxide, can also influence vasodilation. However, this is outside the scope of the current study [9,10]. In this work, we do not explicitly model the mechanisms generating vasomodulatory signals in endfeet. Instead, we approximate their effect on vessel radii using an Ornstein-Uhlenbeck (OU) process.

The exchange of molecules, especially the delivery of essential nutrients, predominantly occurs at the capillary level due to the proximity of capillaries to the tissue. Capillary blood flow increases during neuronal activation. Consequently, alterations in the diameters of blood vessels, especially at the capillary level, significantly modify blood flow in the vascular network. In turn, these changes impact the distribution of vital nutrients essential for supporting neuronal metabolism, including lactate, glucose, and oxygen [11–13].

The astrocytic response shows a noticeable delay of several seconds following neuronal activation as expected from neuronal release of neurotransmitters that initiate astrocytic activation. This response is transmitted to neighboring astrocytes, inducing alterations in the perfusion of nearby capillaries. These changes in perfusion may result from either vasodilation or vasoconstriction in the capillaries [14–17].

Hemodynamic simulations commonly employ mathematical models of cerebral microcirculation that combine biophysical principles with medical imaging data, including X-ray tomographic microscopy [18]. These models integrate anatomical features such as branching patterns and vessel density with hemodynamic and metabolic processes. They provide insights into fundamental anatomical principles, geometric constraints (such as spatial organization, dimensions, and architecture of blood vessels), predictions of cerebral blood flow dynamics, and refined characterizations of metabolite exchange between tissues [19–25]. In addition, established microcirculatory assessment methods, such as laser Doppler flowmetry and capillaroscopy, have laid the groundwork for analyzing vascular function and flow heterogeneity [85], providing context for our modeling approach. In hemodynamic simulations, critical anatomical aspects include the organization of the cerebral microcirculation and structural characteristics such as the arrangement of blood vessels, tissue composition, and spatial relationships between different anatomical elements. Geometric constraints are imposed by the physical geometry of the microvasculature network, including the size, shape, and branching patterns of blood vessels [26,27]. For instance, narrow or tortuous vessels can increase resistance to flow, thereby influencing the overall hemodynamics and affecting distribution of blood and nutrients across different brain regions [28,29]. Astrocytic endfeet may modulate blood flow [30–35] via vascular diameter regulation [36,37].

In this manuscript, we investigate how changes in vessel diameters alter blood flow within a complex vascular network. This study explores the dynamic nature of blood circulation within a complex network of blood vessels, demonstrating how variations in vessel diameter influence overall flow patterns. Additionally, we validate the framework against experimental observations regarding flow values across cortical depth (from cortical layer 1 to layer 6, L1-L6). By employing a multiscale *in silico* model, we integrate the role of astrocytes in regulating cerebral blood flow, a crucial factor in vasodilation and vasoconstriction responses [31]. Our approach provides a foundation for interpreting depth-dependent flow measurements, offering insights that are difficult to obtain through experimental methods alone. This integration highlights the significant contribution of astrocytes to blood flow regulation within the neurovascular unit, thereby establishing a comprehensive framework for understanding cerebrovascular dynamics [38].

## 2. Materials and Methods

Our simulation framework consists of three primary components:

- **Coupling between microvasculature and astrocytic morphologies**. This component ensures that the microvasculature is intricately connected with the synthesized morphologies of astrocytes throughout the circuit.
- **Fluid dynamics model**. This model calculates the flow and pressure within each segment of the vasculature, providing detailed insights into the hemodynamic behavior.
- **Stochastic radii simulation**. We employ a Reflected Ornstein-Uhlenbeck (ROU) process to simulate the dynamic changes in vessel radii, capturing the stochastic nature of vascular adjustments.

Several studies have reported that vasoconstriction is significantly less pronounced than vasodilation across all vessel types [14,39–43]. Thus, we chose to focus solely on vasodilation for the scope of this paper for several reasons. First, vasodilation generally results in larger amplitude changes in vessel diameter compared to vasoconstriction (as for the pial vessels, the amplitude of constriction is significantly lower than for dilation). This makes it easier to detect and measure the effects accurately, providing more reliable data for the model. Second, vasodilation plays a critical role in many physiological and pathological conditions, especially in response to increased metabolic demand and during certain diseases. Therefore, focusing solely on vasodilation enables a more direct examination of these relevant conditions. Lastly, by focusing on only one mechanism, we can simplify the model, reducing computational complexity and allowing for a more targeted analysis.

Algorithm 2.7 also presents a summary of the algorithm for simulating blood flow modulation driven by astrocytic activity.

### 2.1. Digital cerebral microvasculature network

The cerebral microvasculature is an intricate network of interconnected blood vessels [44]. For this study, we use the NGV digital reconstruction of the rat somatosensory cortex (SSCx) presented in [8]. The dimensions of this circuit (954 *µ*m × 1,453 *µ*m ×853 *µ*m) were determined through colocalization of the vascular dataset [8,18] with a neuronal mesocircuit [45]. These dimensions equate to a volume of 1.18 *mm*^3^, which accounts for approximately 0.2% of the rat brain volume. A total of 14,402 astrocytes populate the bounding region, each featuring two endfeet, resulting in a total count of 28,804 endfeet [8] (see Figure 1A, where the astrocytes are blue and blood vessels are red).

**Figure 1.**
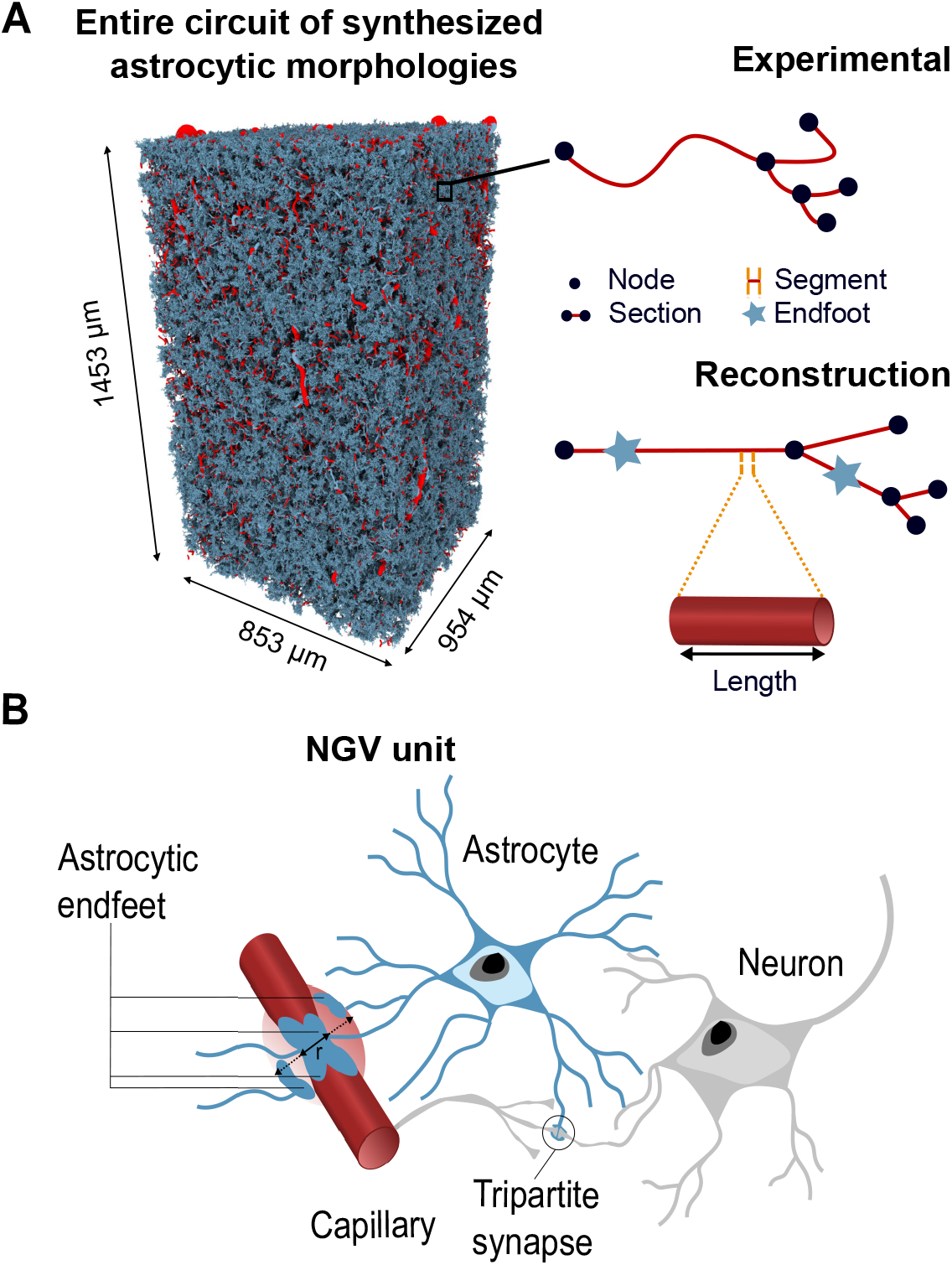
Model overview. (A) The left side shows a complete circuit with synthesized astrocytic morphologies developed by Zisis et al. [8]. Astrocytes are shown in blue and blood vessels in red. On the right there is a schematic depiction of a realistic microvascular network sample, comprised of nodes and edges. (B) NGV unit. Neurons are depicted in gray, astrocytes in blue, and capillaries in red. Astrocytes contact synapses, wrap around them, and extend their perivascular projections to the surface of blood vessels, where they form endfeet. *r* stands for radius.

The vasculature dataset is represented as a graph composed of nodes and edges. Each node is characterized by its coordinate position and its radius, while edges indicate the connections between nodes. The length of an edge is simply the distance between the connected nodes, and the edge radius is defined as the arithmetic average between the node radii. The microvascular network (see Figure 1) serves as the structural framework for the placement of astrocytic endfeet.

In a realistic network each edge corresponds to a blood vessel, it is located between two bifurcations (branching nodes), and typically exhibits tortuosity. In the model we simplified the network reconstruction by disregarding the tortuosity while still accounting for the actual lengths of the tortuous vessels. As a result, each edge was modeled as a straight pipe with a fixed radius. Figure 1A illustrates a representation of a cerebral microvascular network.

### Volumetric analysis of cortical layers and vascular system

Experimental data of the vessel volumes within each cortical layer. CAP stands for capillaries, while LV stands for large vessels. The low numbers within L6 in all columns can be attributed to the limitations of the experimental dataset. The flow ratio is the proportion of flow inside capillaries and large vessels. The values do not add up to 100% because we do not consider middle size vessels.

The vasculature volume was extracted from the rat SSCx [18]. It was then aligned with the cortical column microcircuit using cortical layer boundaries as the neuronal region in the rat SSCx was usually subdivided in six layers [46]. A significant section of the vasculature was located above L1 and needed to be taken into account as it contains the main arteries. Moreover, the current vasculature volume [18] is truncated at approximately one-fourth of the height of L6, from the top. Therefore, three-fourths of L6 down to the white matter have no vasculature in this model. Thus, vasculature and microcircuit volumes do not perfectly overlap (see Figure A5).

Table 1 provides data of several quantities inside the six-layered structure such as the volume of the layer, the vascular volume, the proportion of the total volume occupied by blood vessels, and the proportion of flow passing through capillaries and large vessels.

**Table 1.** Vasculature network parameters.

| Layer | Layer volume ($\mu\text{m}^3$) | Vascular system volume ($\mu\text{m}^3$) | Volume occupied by vessels (%) | Flow ratio (%) | |
| --- | --- | --- | --- | --- | --- |
|  |  |  |  | CAP | LV |
| L1 | $1.29 \cdot 10^8$ | $5.03 \cdot 10^6$ | 3.88 | 66.17 | 3.27 |
| L2 | $1.16 \cdot 10^8$ | $1.16 \cdot 10^6$ | 4.24 | 67.25 | 2.64 |
| L3 | $2.77 \cdot 10^8$ | $9.39 \cdot 10^6$ | 3.40 | 62.98 | 2.76 |
| L4 | $1.48 \cdot 10^8$ | $4.35 \cdot 10^6$ | 2.92 | 26.61 | 1.28 |
| L5 | $4.12 \cdot 10^8$ | $8.25 \cdot 10^6$ | 2.00 | 35.46 | 1.33 |
| L6 | $5.49 \cdot 10^8$ | 3.47 | 0.06 | 53.15 | 0 |

The low percentage of vessels within L6 can be attributed to the limitations of the experimental dataset. This dataset only contains a cropped portion of L6, resulting in a relatively small representation of this layer. It was imperative to consider L6 while conducting our analyses, even if its dataset was limited. The flow ratio highlights the proportion of flow in capillaries and large vessels. The sum is smaller than 100% as we exclusively considered capillaries (diameters smaller than 6 *µm*) and large vessels (diameters larger than 14 *µm*) among various types of vessels. The decision to exclude other vessels was made to emphasize the distinctive behaviors of capillaries and large vessels. Capillaries are crucial for nutrient and gas exchange at the cellular level, while large vessels are vital for rapid blood transport and pressure regulation. By focusing on these extremes, our analysis can better capture the specific roles and responses of these vessel types under various physiological and pathological conditions. This approach facilitated easier validation of our results against the existing literature because extensive research and data are more readily available for these vessel types, providing well-documented benchmarks for comparison [47–49].

### 2.2. Mathematical framework of the blood flow model

Here, we present the mathematical framework of the fluid dynamics model that characterizes the flow and pressure of blood within the vascular system. The vasculature was modeled as an intricate graph of interconnected blood vessels. In turn, each one of them was represented as a tube of length *l* and with circular cross section *A*(*t*) = *π r*(*t*)^2^, where *r*(*t*) is the radius and *t* is time. The vessel section *A*(*t*) could change over time due to the effect of endfeet and was constant along the longitudinal axis. The blood was treated as an incompressible and viscous fluid circulating within each vessel. Blood velocity *u*(*t, x*) and pressure *p*(*t, x*) were assumed to be constant in the radial direction. Therefore, they only depend on the longitudinal spatial dimension *x*, and the entire system could be considered one-dimensional.

The mathematical framework considers the following 1D Navier-Stokes momentum equation:

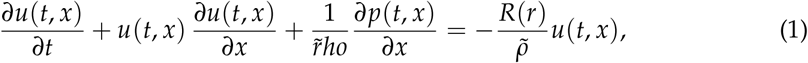

where 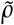 is the constant fluid density, and *R*(*r*) is the resistance with the following expres-sion:

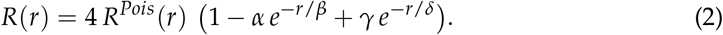

*R*^*Pois*^ (*r*) is the classical Poiseuille resistance defined as:

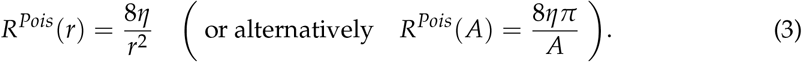

The radius is expressed in (*µ*m) while *η* represents the viscosity of the plasma (1.2 ·10^−6^ *g* · *µm*^−1^ · *s*^−1^). In this study, the blood flow was modeled assuming a constant plasma viscosity. While it is well-established that blood exhibits non-Newtonian behavior, with viscosity that varies as a function of shear rate and hematocrit concentration, this simplification allows for a more tractable analysis of vascular flow dynamics, particularly in complex networks. Previous studies have demonstrated that such an approach can still yield reasonably accurate insights when the primary focus is on comparative or qualitative trends rather than precise rheological detail [72].

The resistance in Eq (2) was a modification introduced in [44] of the classical Poiseuille resistance to account for the influence of hematocrit. We used the same values of the model parameters *α* = 0.863, *β* = 14.3*µm, γ* = 27.5, and *δ* = 0.351*µm* provided in [44].

Making the further assumption that the speed was constant with respect to time and space i.e. 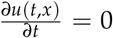 and 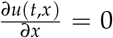, it follows that Eq (3) becomes the well known

Hagen-Poiseuille equation:

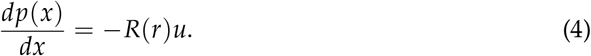

Let us consider a single edge between node *i* and node *j*. We denoted with *R*_*ij*_ the resistance *R*(*r*) associated with the selected edge, and with *A*_*ij*_ the section area. Let us integrate both sides of the equation between the positions *x*_*i*_ and *x*_*j*_ of the two nodes:

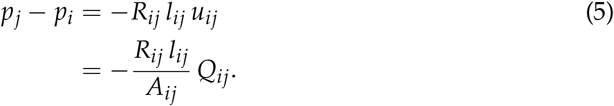

The term *u*_*ij*_ indicates the velocity between *i* and *j, l*_*ij*_ is the length of the edge, and *Q*_*ij*_ = *u*_*ij*_ *A*_*ij*_ is the flux (also called flow in the next sections). Let us call:

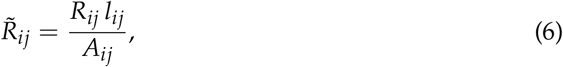

such that

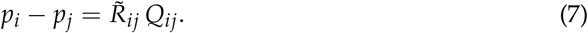

We considered the following flux conservation condition, and replaced the expression of *Q*_*ij*_:

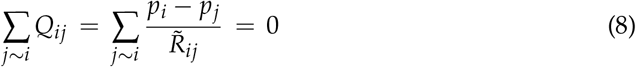

where the sum was over nodes *j* connected to node *i*. It was convenient to rewrite our equations into matrix-vector notation. Let us introduce the *N × M* oriented incidence matrix *B* (with *N* nodes and *M* edges) defined as *B*_*i,k*_ = 1 and *B*_*j,k*_ = −1, with *k* representing the index of the oriented edge connecting nodes *i* to node *j*. The signs indicate the orientation of the graph edges. We defined the graph Laplacian *L* as

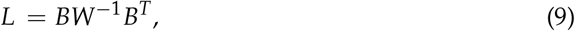

where *W* is the *M× M* diagonal matrix of edge resistances (i.e.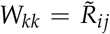). We rewrote Eq (8) into matrix-vector notation, including the boundary conditions, as:

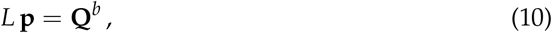

where **p** is the unknown pressure vector and **Q**^*b*^ is a given vector containing the boundary flows on the nodes of degree one, and with values equal to zero on the internal nodes with degree greater than one. The vector 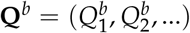 satisfies the property 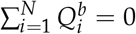, corresponding to the vanishing total flow condition.

Once the pressure is computed, we could retrieve the flux vector since Eq (7) can be written in vector form as:

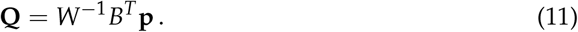

Often the boundary flow is approximated using various methods such as building a larger homogeneous vasculature with simple boundary conditions [50].

Section 2.3 explains how we computed the boundary flows. Once they were set, the solution of Eq (10) could be computed up to a constant pressure value. It is common practice to set it equal to the external pressure: 10 *mmHg* (1.33 *g* · *µm*^−1^ · *s*^−2^) [51].

### 2.3. Blood flow on the boundary nodes

The static flow model described in Section 2.2 computes the flow along each edge within the graph, given the flows assigned at the boundary nodes. The simulation time [0, *T*] was discretized in ℐ time steps and ℐ+ 1 time points. We solve Eq (10) and Eq (11) at each time point with different boundary conditions and different values of vessel radii. The present section focuses on the time-dependent boundary conditions **Q**^*b*^ (*t*) for *t* ∈ [0, *T*] while Section 2.4 shows how radii dynamics are modeled.

Our approach has three main components:

1. **Entry nodes selection**. The nodes where blood is introduced into the system.
2. **Formulation of a time-dependent flow model**. It is needed to characterize the injected flow within the entry nodes.
3. **Exit flow calculation**. Computation of the quantity of blood leaving the vasculature through the exit nodes.

First, we injected the flow into the largest *ñ* nodes (with the greatest diameter) located on the top region of the vasculature. This choice is consistent with the actual artery location and flow direction measured in the rat SSCx [52]. In particular, we defined the top as the zone where: *y* > *y*_*min*_ + 0.95 (*y*_*max*_ − *y*_*min*_) being *y* the vertical coordinate of a cortical column oriented with the lowest layers on top (L1), down to the lowest layer (L6) at the bottom. This is consistent with the actual artery location and flow direction measured in the rat SSCx [52].

After that, we needed to determine the amount of flow to inject. [52] provides the blood velocity time series for large vessels. Figure A**??** shows that the velocity signal has a sinusoidal shape. It was convenient to fit the empirical velocity signal with a sine wave:

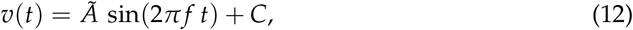

where *Ã* ∈ ℝ is the sine amplitude, *f* ∈ ℕ is the sine frequency and *C* ∈ ℝ is the signal baseline. Considering a unit interval for the time, *t* ∈ [0, 1], such that the sine domain is a multiple of a 2*π* pe<u>r</u>iod, the parameters were estimated as follows: *C* is the mean value of the signal, *Ã* is 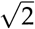 times the standard deviation of the signal, and *f* is the number of peaks. Following this approach, the fitted parameters are: *Ã* = 6119 *µm*· *s*^−1^, *f* = 8 *s*^−1^, and *C* = 35000 *µm*· *s*^−1^.

Once we know the velocity *v*(*t*) we computed the flow to inject into the entry node *i*:

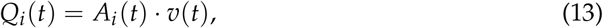

where *A*_*i*_(*t*) is the area of the entry node *i* at time *t*.

The last step was to determine the flow in the exit nodes. Let us suppose that there are 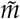 exit nodes. We defined the weight *w*_*j*_(*t*) as:

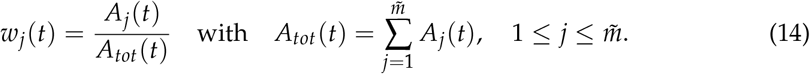

From the total flux conservation law, we know that the total exit flow 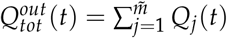 and the total entry flow 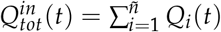 must satisfy:

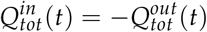

at each time *t*∈ [0, *T*].

Thus, the flow in the exit node *j* is:

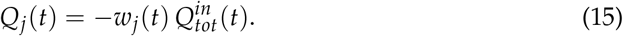

Section A.1 provides a detailed overview of the history of exit flow computation, leading to the current version discussed above.

### 2.4. Stochastic endfeet activity

As explained in Section 2, our study focuses exclusively on vasodilation. Consequently, we employ a Reflected Ornstein-Uhlenbeck (ROU) process [53,54] with a reflection barrier at zero to prevent negative values, thereby accurately replicating the vessel radii dynamics influenced by endfeet activity.

A generic ROU process *X* = {*X*_*t*_ : 0 ≤ *t* ≤ *T*} is described by the following stochastic differential equation:

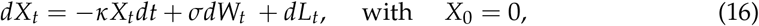

where *κ* is the reversion to zero coefficient and *σ* is the diffusion coefficient. *W* = *{W*_*t*_ : 0 ≤ *t* ≤ *T}* is a Wiener process, and *L* = *{L*_*t*_ : 0 ≤ *t* ≤ *T}* is the minimal non-decreasing process which makes *X*_*t*_ ≥ 0 for all *t* ≥ 0, also known as the local time.

The simulation time [0, *T*] is discretized in ℐ ∈ ℕ time steps of size Δ*t*. Let us indicate with *X*_*i*_ the value of *X* at the time step *i* ≤ ℐ. *X*_*i*_ can be calculated using the recursive relation:

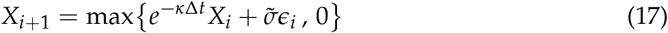

where 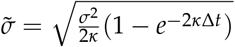 and *ϵ*_*i*_ ∼ 𝒩 (0, 1) is a standard Normal random variable.

The radius dynamics *r*_*i*_ for 0 ≤ *i* ≤ ℐ could be modeled as

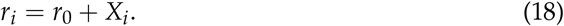

where *r*_0_ is the reference resting state radius.

### 2.5. ROU calibration

The ROU parameters *κ* and *σ* are calibrated from the values of the radius maximal extension *r*_*max*_ and the mean time *t*_*max*_ to reach it from *r*_0_. It is useful to recall that the asymptotic distribution of the ROU process follows the half-Normal law:

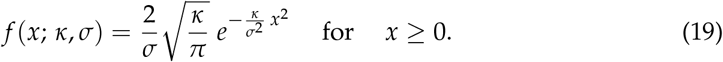

In the region *x* ≥ 0, this distribution is exactly two times the asymptotic distribution of the OU model, which is a normal distribution with mean 0 and standard deviation 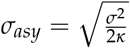.

For convenience, we decided to measure the distance from the origin as a multiple of *σ*_*asy*_. Let us introduce the new parameter *c* > 0 and set:

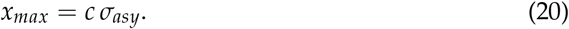

Our main modeling choice is to set *c* = 2.8. With this assumption, we want to introduce a realistic maximum value for the ROU process. The probability ℙ(*X*_*T*_ > *x*_*max*_) ≈ 0.5% is small enough to ignore. Larger values of *c* would make it difficult for the process to reach *x*_*max*_. In Figure A**??** we can see a sample path and an histogram of the ROU process with the new introduced maximum value.

In the calibration process, we can retrieve the value of *x*_*max*_ from the experimental values of *r*_*max*_ and *r*_0_ by setting *x*_*max*_ = *r*_*max*_ − *r*_0_. Eq (20) expresses the parameter *σ* in terms of *κ*:

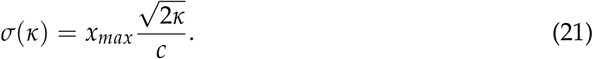

The expected time *u*(*x*) to reach *x*_*max*_ for a ROU process starting from *x* is the solution of the following differential equation (see [53]):

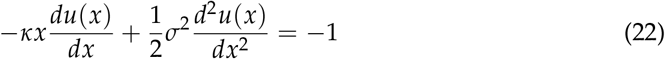

with *u*^*′*^ (0) = 0 and *u*(*x*_*max*_) = 0. This boundary value problem has mixed boundary conditions. We numerically integrate this second-order, ordinary differential equation employing a finite difference scheme using a backward difference for the first-order derivative and a central difference for the second-order derivative.

The expected time starting from *x* = 0, obtained from Eq (22), depends only on the parameter *κ* since *σ* can be substituted from Eq (21). For this reason, we indicate this dependence as *u*_*κ*_(0):

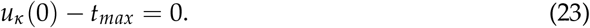

This equation can be solved numerically for *κ* using a root-finding algorithm, such as the Brent method [55].

### 2.6. Passive response of the network

In our model, the phase corresponding to the passive response of the vasculature corresponds to the phase where when there is no astrocytic activity (i.e. no noise). We can remove the Wiener noise in Eq. (16) by setting *σ* = 0.

Let us define *τ* as the time at which the endfeet activity ceases. During the passive response phase, for *t* > *τ*, the radius dynamics decays exponentially as:

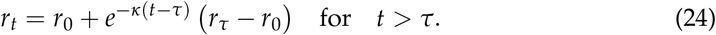

The average radius dynamics among all edges, 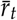, can be determined through:

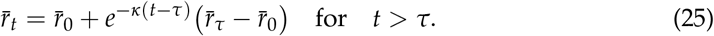

Here 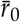 and 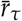 represents the average blood vessel radius at time 0 and *τ* respectively.

### 2.7. Algorithm implementation

In the previous sections, we introduced a mathematical model for computing flow and pressure within a vasculature in presence of astrocytic activity. Algorithm 1 outlines all the necessary steps for the simulation.

#### Algorithm 1

Simulation of blood pressure and flow during astrocytic stimulation.

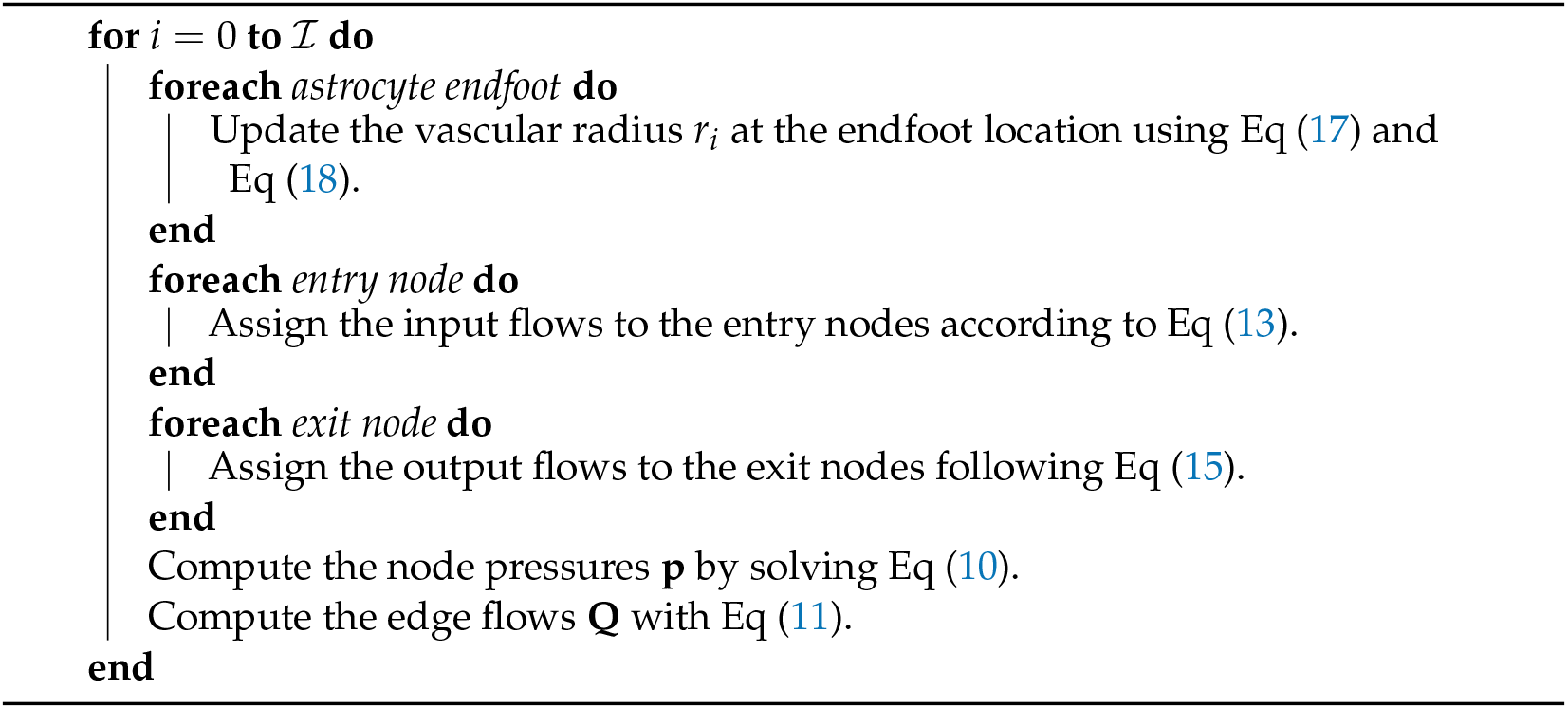

The algorithm, implemented in Python, was released as an open-source package named AstroVascPy. The source code is available for public access on GitHub [56]. The software was designed for scalability across diverse vascular network datasets, with the possibility to use parallel computation on modern HPC clusters.

The part of the code that requires the most computational resources is the solution of the sparse linear system Eq (10). After several tests, we found that for graphs with ∼ 10^6^ nodes it is convenient to solve the system with a direct method, while for very large graphs (more than 10^7^ nodes) it is more efficient to use an iterative parallelized algorithm. For this purpose, we relied on the Python package *petsc4py* [57], PETSc python binding, a well known C library for numerical parallel computation.

The particular vasculature network considered in this paper has *N* = 1, 351, 448 nodes and *M* = 1, 349, 411 edges. In our analyses, we reduced the graph to its largest connected components. We then got *N* = 1, 049, 221 and *M* = 1, 051, 911. The Laplacian *L* in Eq 10 is a sparse *N ×N* matrix. In this case, the system is quite small and could be solved quickly by a direct method such as the LU decomposition. We relied on the Python library *scipy.sparse* for this computation.

### 2.8. Mathematical framework for quantitative analysis

In this section, we define key quantities that will be essential for analyzing the results in the Section 3 Section. At each time step *i*, and at each edge *k* we indicate the flow as *Q*(*i, k*) and the radius *r*(*i, k*). For the edges connected to endfeet, quantities are denoted with the superscript *e*, e.g. *Q*^*e*^ (*i, k*), *r*^*e*^ (*i, k*), etc.

We define the *Resting State ratio* for the flow and the radius as follows:

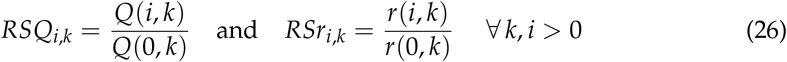

Let us consider a subset of the set of all edges that contains *M*^*′*^ edges, with *M*^*′*^≤ *M*. We define the *Average Ratio* for the flow, over a selected subset of edges as:

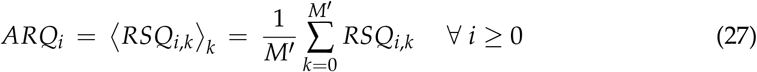

We use ⟨ ⟩ brackets to indicate an arithmetic average with respect the variable indicated in the subscript.

When we take an average over a set of edges and over time, we indicate it as

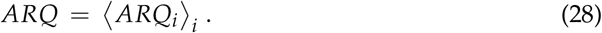

#### 2.8.1. Order ratios

The distance order *m* of an edge 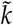 with respect to target edge *k* is defined as the minimum number of connected edges we need to traverse to reach 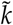 starting from *k*. For example, if two edges share a node, their distance order is 1. If you need to pass by an extra node to move from the initial edge to the final edge the distance order is 2 and so on.

Now, let us introduce the concept of *Order Ratio* for the flow, to quantify how the astrocytic activity on an edge *k* affects segments in the surrounding vascular network. We define it as:

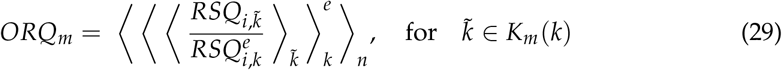

where *K*_*m*_(*k*) is the set of all edges of distance order *m* from a target edge *k*. The most interior average was done over the elements of *K*_*m*_(*k*). The average in the middle was done over all edges *k* connected with endfeet. And finally, the last average was done over all time points *n*.

## 3. Results

In this paper, our primary goal was to characterize the interactions between astrocytic activity and blood flow within a large and complex vascular network. To achieve this, we developed a model that simulates the effect of astrocytes on vessel radii using a reflected Ornstein-Uhlenbeck process (see Section 2.4). Subsequently, we employed the 1D Navier-Stokes momentum equation to compute blood-flow in each segment of a given vascular network (see Section 2.2). We decided to apply this model to the rat SSCx vasculature from Reichold et al. [18] since astrocyte placement was previously performed on this vasculature [8].

### 3.1. Model presentation

Throughout the paper, we focused our analyses on the six cortical layers of the rat SSCx due to their distinct functional and structural characteristics (see Figure 2A). Understanding the organization and dynamics of these layers is crucial for capturing the complex neural processes underlying sensory perception and integration [58–60]. Traditional vascular studies typically categorize vessels into two types: capillaries with a diameter *d* [4 − 6] *µm* and larger vessels (*d* ≥ 14 *µm*). We thus used a similar categorization to study the interaction between endfeet activity and the vascular network. The astrocytic endfeet distribution is quite constant across cortical layers one to four with around 800 endfeet per bin of 25 *µm* along the cortical depth (Figure 2B). The number of endfeet then decreases within L5 from 800 to 500 endfeet per 25 *µm* bin. On the other hand, we observed a prevalence of capillaries within the deeper layers (L4-L5), while large vessels are prevalent in upper layers (L1-L2), see Figure 2C. As expected, the number of large vessels decreased in deeper layers. Vasculature dataset incompleteness did not allow us to observe the endfeet distribution in L6. A typical simulation consisted of a three-second long injection of flow into the vasculature entry nodes (Figure 2D, gray rectangle), accompanied by a stochastic expansion of the radii of the vasculature sections connected to the endfeet (see Section 2). For the sake of simplicity, in the following analyses, we use a constant blood velocity of 3.5 10^4^*µm* · *s*^−1^. This is equivalent to set *Ã* = 0 in Eq (12). We then simulated the next two seconds with no astrocytic activity to observe the vasculature response through a relaxation period (see Section D). Thus, we computed the flow and pressure variations at each time step across all the vasculature segments during the time in which the astrocytes are active (from zero to three seconds) and the relaxation period (from three to five seconds). The results can then be easily classified based on various criteria, such as cortical layers (L1 to L6) and vessel types (capillaries or large vessels). Figure 2D-F show typical simulation results. Figure 2D shows time series of mean flow rate grouped per layer and type of vessels (large vessels on top and capillaries on bottom). We observed across type of vessels and endfeet activity states, the mean flow consistently exhibits higher values in cortical layers L1 and L2, compared to deeper layers (see Figure 2D). We observed a very similar mean flow dynamics in both capillaries (Figure 2D, on bottom) and large vessels (Figure 2D, on top). There was a difference of two orders of magnitude between the mean flow values of capillaries and large vessels (10^5^ and 10^7^, respectively). The mean blood flow dynamics in capillaries (Figure 2D, on bottom) tends to form two groups: L1 and L2 on one hand, and L3-L6 on the other hand. This highlights the difference between the first two layers and the others about the quantity of flow. Similarly, the mean flow in large vessels (see Figure 2D, top), also shows similar behavior in L1 and L2, and progressively decreases in deeper layers.

**Figure 2.**
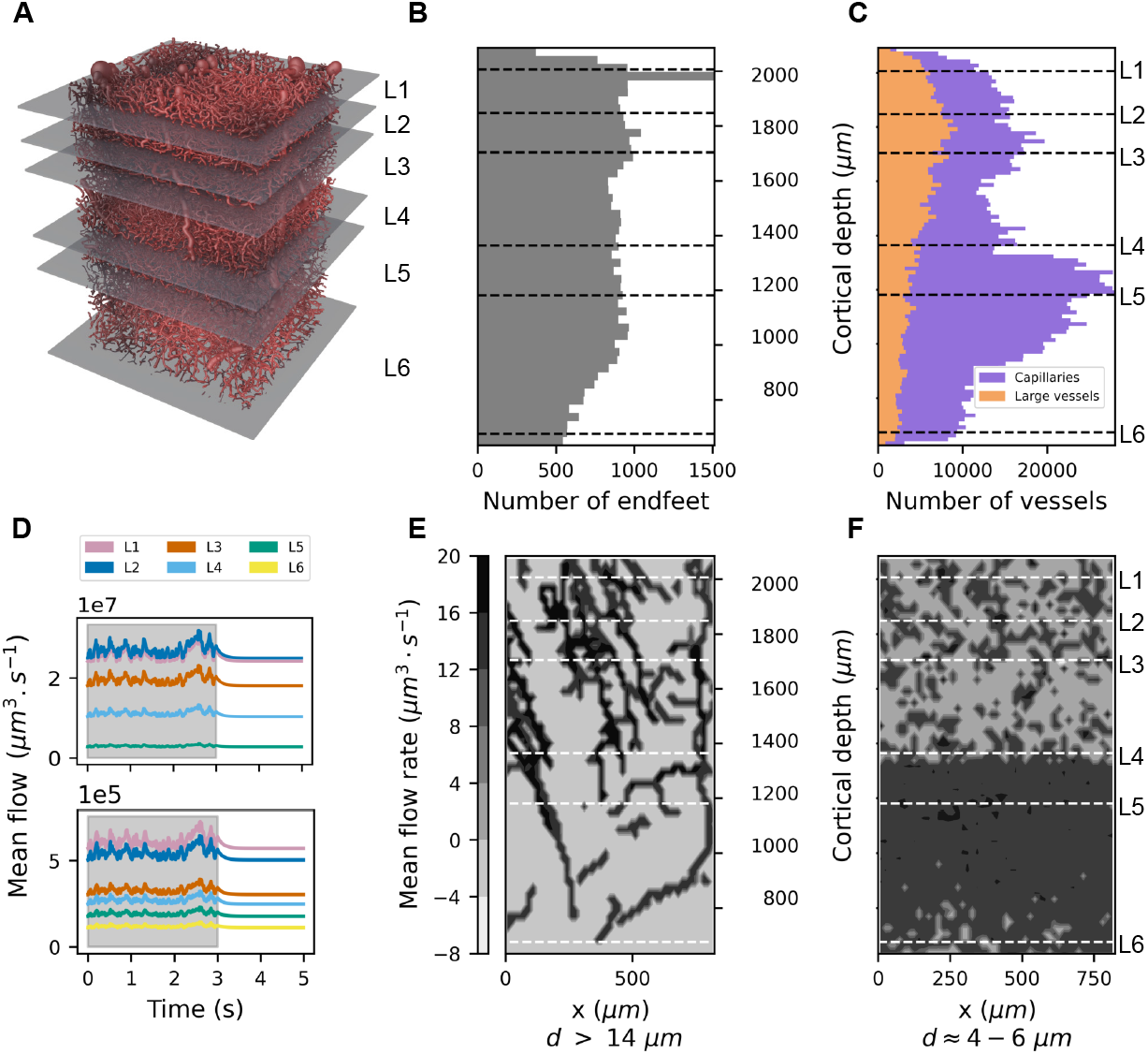
Model presentation. (A) Vasculature segmented into six cortical layers in the rat SSCx separated by gray planes. (B) Endfeet distribution along the cortical depth in the vasculature [8]. (C) Spatial distribution of capillaries and large vessels along the cortical depth [8]. (D) Time series illustrating the flow variations averaged across all large vessels (top) and capillaries (bottom) in each cortical layer (color-coded) in response to a three-second astrocytic stimulation (gray rectangle). Since there are no large vessels in L6, a corresponding time series is missing from the legend. (E) Heatmap of the mean flow in the vasculature at a specific time (*t* = 3 *s*) within the simulation, calculated as an average over a surface area of 17 *µm ×* 21 *µm* within large vessels (diameter *d* ≥ 14 *µm*). (F) Heatmap of the mean flow in the vasculature at a specific time (*t* = 3 *s*) within the simulation, calculated as an average over a surface area of 17 *µm ×* 21 *µm* within capillaries (diameter *d* ≈ 4 − 6*µm*). The vertical axis in B-C-E-F represents the cortical depth.

Figure 2E and Figure 2F depict heatmaps of the mean flow in the vasculature at a specific time (*t* = 3 *s*) for large vessels and capillaries respectively. The cortical depth is represented by the y-axis while the x-axis represents a 2^*nd*^ dimension of the vasculature. The values of the mean flow were then concatenated along the z-axis to create the heatmaps representation. In the deeper cortical layers, there are more capillaries, resulting in more blood and higher total blood flow (Figure 2F). Conversely, the upper layers have more large vessels, leading to increased blood volume and higher total blood flow in that region (Figure 2E).

Overall, these results suggest that most of the variance of flow during the passive phase is attributed to the architecture of the vasculature network. The distribution of blood flow was categorized into supragranular and subgranular layers and analyzed throughout the paper.

### 3.2. Astrocytic activity affects flow in the vascular network

To characterize the model, we quantified the changes in vessel radii due to astrocytic activity and the subsequent alterations in the blood flow.

Responses to endfeet activity differ between capillaries and large vessels (radii modulation in Section 2). We used average data reported in the literature to set the maximum dilation, the *maximum radius ratio*, for both vessel types. In particular, it was set to 1.38 for capillaries and 1.23 for large vessels, based on [14,39,41–43,61–63]. We also set the time it takes to reach the maximum radius, the *time to peak*, for both vessel types. It represents the average duration from the onset of the stimulus to the time where maximal dilation is reached. In our simulations, we fixed this *time to peak* at 2.7 s for capillaries and 3.3 s for large vessels (see Section 2, Table 2 and Table 3, and [14,39,41–43,61–63]). Our choice of a 2.7 s time scale for the hemodynamic response represents an averaged simplification that captures the general dynamics observed experimentally. We acknowledge that fast (hundreds of milliseconds to seconds) responses are thought to be driven by rapid potassium signaling [79], while slower modulation occurs on the order of tens to hundreds of seconds, influenced by neuromodulators [78,83]. Nevertheless, the 2.7 s time scale was chosen to balance these effects and reflect intermediate responses typical in neurovascular modeling. In Figure 3A-B, we highlight the distribution of the resting state ratio for radius and flow. The histograms in Figure 3A-B consider all the edges connected exclusively to a single endfoot, at every time point. In accordance with the literature ([14,39,41–43,61–63]) and our model implementation (see Section 2), 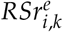 (see 26) is concentrated between 1 and 1.3, showing a peak close to 1.0 and gradually decreasing until 1.3.

**Table 2.** Simulation parameters in capillaries.

| Number of time steps | Time step size | Time to peak | Maximum radius ratio |
| --- | --- | --- | --- |
| 500 | 0.01 s | 2.7 s* | 1.38* |

Values marked with an asterisk (\*) are based on the literature data [14,39,41–43,61–63].

**Table 3.** Simulation parameters in large vessels.

| Number of time steps | Time step size | Time to peak | Maximum radius ratio |
| --- | --- | --- | --- |
| 500 | 0.01 s | 3.3 s* | 1.23* |

Values marked with an asterisk (\*) are based on the literature data [14,39,41–43,61–63].

**Figure 3.**
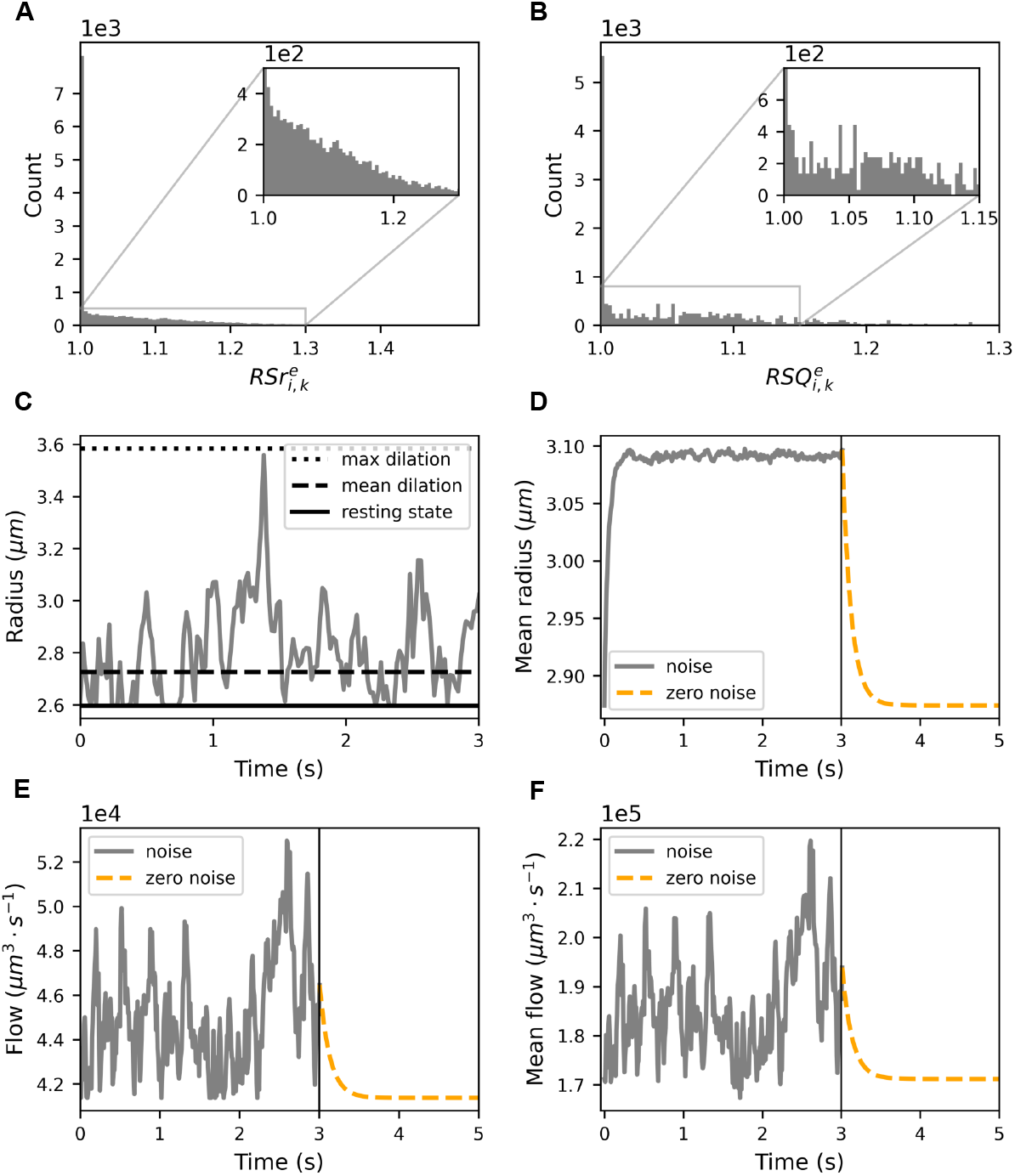
Dynamic analysis of blood vessel radii and flow in response to astrocytic activity. (A) Histograms showing the distribution of 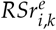 and (B) 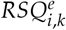 for each edge connected to only one endfoot and for each time point (see Section 2.8). (C) Time series plot showing the radius dynamics for a single segment. The solid black line represents the resting state radius, the dashed black line represents the average radius over time, and the dotted black line indicates the theoretical maximal extension of the radius. (D) Time evolution of the mean radius across all blood vessels when stimulation is halted after three seconds. (E) and (F) present the equivalent time series for the flow. (D-F) The solid gray line represents the stimulation period, while the dashed orange line illustrates the passive phase, after the stimulus ceased.

Similarly, in Figure 3B the distribution of the resting state flow ratio captures the temporal changes in flow dynamics of the vessel segments connected to endfeet. In other words, while in Figure 3A we explored variations of the input (i.e. the radii 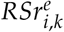), in Figure 3B we focused on the model output (i.e. the flow 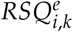 and 1.15 presents a high concentration of peaks of 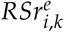). The area between 1 while the distribution remains relatively flat. The irregular shape of the flow distribution indicates a highly nonlinear interaction between the radius dynamics (input) and the observed flow (output).

Such near-unity flow ratios indicates a predominant occurrence of minimal changes in flow between successive time points. This observation underscored the robustness of the neurovascular system in maintaining cerebral blood flow within a narrow range, even in the presence of dynamic physiological processes such as neuronal activity and metabolic demands [64–66].

Figure 3C presents the path of the simulated dynamics for a single capillary taken in the center of the vasculature. The radius follows Eq (18). The maximum dilation is obtained from the literature data. Table 2 and Table 3 outline all the additional parameters used in the simulation.

Figure 3D shows the evolution of the mean radius during the simulation. Figure 3E presents the time series of the flow inside the capillary considered in Figure 3C.

Figure 3D-F presents the exponential decay profile of the mean radius, as predicted in Section D. According to Eq (A4), the typical timescale for the vasculature to return to the resting state, the characteristic time, is approximately 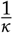.

Figure 3D shows the mean of all radii during simulated astrocytic stimulation over time. An increase in astrocytic activity led to a temporary increase in the mean value of the radii across the vasculature, indicative of an average expansion of the total vasculature volume.

Similarly, the equivalent plot for the flow in Figure 3E illustrates the response of a single blood vessel. Here, the increase in astrocytic activity leads to a transient increase in flow.

We show in Figure 3F the time evolution of mean flow across all blood vessels. As in the previous cases, it presents higher noisy values during the initial stimulation phase, and then it decreases exponentially in the following passive phase. The reason for a noisy path of the mean flow is due to the stochastic nature of the flow we inject into the vasculature. We keep the blood speed constant and multiply it by the sectional area of the three entry nodes, which are all connected to the endfeet.

The similarity between the curves in Figure 3E and Figure 3F could be explained by the principles of vascular physiology and the dynamics of NGV coupling, i.e. cerebral autoregulation and flow-metabolism coupling [105,106].

### Simulation results reproduce flow ranges in the literature

To validate the model, we compared its outputs with flow rates and velocities reported in the literature.

In our simulations we imposed the value of the flow **Q**^*b*^ on the boundary nodes Eq (10). Thus, the validation focused solely on the blood vessels in the internal part of the vasculature, in the region where: *x* ∈ [67.7, 622.3] *µm, y* ∈ [1033.5, 1684.9] *µm*, and *z* ∈ [273.27, 926.73] *µm*. It yields a 17% of the vasculature falling within the defined region. In this way, we minimized the possible distortions introduced by the boundary conditions and took into account contributions from all nodes in the vasculature.

The validation of the simulation in Table 4 and Section 3.2 focused on an analysis of flow values in capillaries categorized by their diameters.

**Table 4.** Refined validation of the simulation outputs. The flow values in capillaries are grouped according to their diameter.

| Diameter ($\mu\text{m}$) | Number of edges | Flow in capillaries ($\mu\text{m}^3 \cdot \text{s}^{-1}$) | | |
| --- | --- | --- | --- | --- |
|  |  | min | max | mean |
| 1-2 | 7,106 | 0.01 | $2.22 \cdot 10^5$ | $6.95 \cdot 10^3$ |
| 2-3 | 21,140 | 0.11 | $2.72 \cdot 10^6$ | $3.03 \cdot 10^4$ |
| 3-4 | 14,228 | 1.60 | $5.52 \cdot 10^6$ | $1.60 \cdot 10^5$ |
| 4-5 | 90,748 | 7.74 | $9.05 \cdot 10^6$ | $4.40 \cdot 10^5$ |
| 5-6 | 59,670 | 0.79 | $1.89 \cdot 10^7$ | $9.10 \cdot 10^5$ |
| 6-7 | 24,853 | 8.83 | $2.14 \cdot 10^7$ | $1.48 \cdot 10^6$ |

In the literature, it is common practice to express flow rates in large vessels in 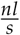 and flow rate in capillaries in terms of the number of red blood cells (RBC) per second (*RBC*· *s*^−1^). Table A3 recaps unit conversions and the parameters used in the simulations.

When interpreting flow values from the literature, it is crucial to consider potential selection biases and undetected artifacts. Biases may arise from the selective measurement of vessel segments, which might not be representative of the entire vascular network. Furthermore, blood flow measurements confined to specific regions or under specific conditions may not fully capture the variability present throughout the vasculature. Moreover, artifacts can stem from the resolution limits of instruments, leading to discrepancies between measured and actual values.

We classified capillaries into six categories, each distinguished by a diameter increment of 1 *µm* (ranging from one to seven micrometers), to closely align with existing literature [50]. Minimum, maximum, and mean values were recorded for each segment throughout the simulation. Next, we averaged these values over all the segments of a particular diameter range. Table 4 shows the relationship between capillary diameter and flow dynamics within the rat SSCx. Our analysis confirmed an increase in flow rates as capillary diameter increases, with the highest flow observed in capillaries within the 6-7 *µm* diameter range.

The flows and velocities produced by the model align with the literature data for both capillaries and large vessels. In cases where average data was missing in the literature, only a range of values was reported from Shih et al.[69]. Flow and velocity ranges from the model were wider than the ones from the literature (see Figure 4A and Figure 4B). It is worth noting that our validation encompasses data from thousands of vessels, a scale larger than in the typical literature, where data often comes from only tens or hundreds of vessel segments. Therefore, it is expected that our results exhibit a broader range. Nevertheless, the bulk part of the model data (2^*nd*^ and 3^*rd*^ quartiles of the box-plots) were in the typical literature range close to 10^4^ *µm*^3^ · *s*^−1^ for the flow and 10^4^ *µm* · *s*^−1^ for the velocity. This was further supported by the detailed distribution profiles of the flow values (see Figure 4C and 4E) and velocity values (see Figure 4D and Figure 4F) for capillaries and large vessels respectively. Table A1 and Table A2 illustrate validation flow and velocity.

**Figure 4.**
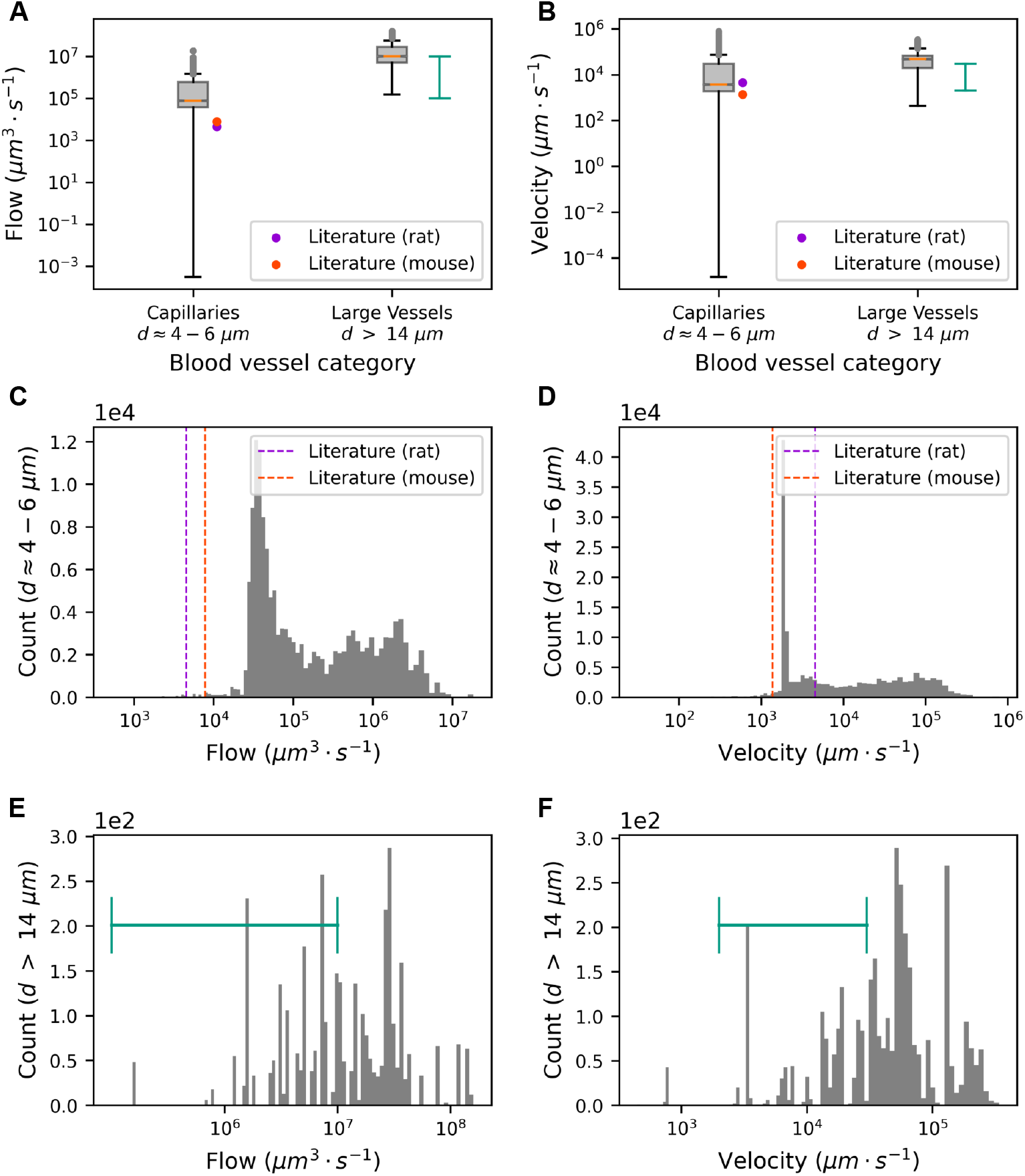
Validation of our simulation results against the existing literature data. Blood flows (A) and velocities (B) are evaluated for both capillaries and large vessels. Mean values of simulated flow/velocity are produced by averaging over all segments throughout the active astrocytic phase. Their distribution is shown in the boxplots. Gray dots show outliers, orange lines indicate the median flow in blood vessels. Dot markers represent values from previous studies [67,68], color-coded by species. Error bars in turquoise green depict flow and velocity minimum and maximum observed for rats in [69]). Distribution of blood flow (C) and velocity (D) in capillaries (diameter d ≈ 4 − 6 *µm*) plotted on a logarithmic scale. The dashed lines indicate the literature values for rat and mouse. Distribution of blood flow (E) and velocity (F) in large vessels (*d* ≥ 14*µm*) plotted on a logarithmic scale. Error bars in turquoise green depict flow and velocity ranges observed for rats in [69].

### Global architecture of the vasculature affects astrocytic activity

In Figure 5, we compared the extent of flow changes carried out by capillaries versus large vessels. Interestingly, the total volume of capillaries was 1.24 · 10^7^*µm*^3^, occupying 35% of the vascular surface, while the volume of large vessels was 1.0 · 10^7^*µm*^3^, occupying 28% of the vascular volume. As expected, Figure 5A shows that average flow ratios across all vessel types exhibit a similar order of magnitude. Smaller vessels have a higher average resting state flow ratio. Figure 5B, instead, examines the distribution of the flow ratio across layers. The three types of vessels appeared to have the same proportion (one third) across all layers except L5. Overall, capillaries, large vessels, and other vessels constituted 35.05%, 31.95%, and 33.00% of the total resting state flow ratio distribution, respectively.

**Figure 5.**
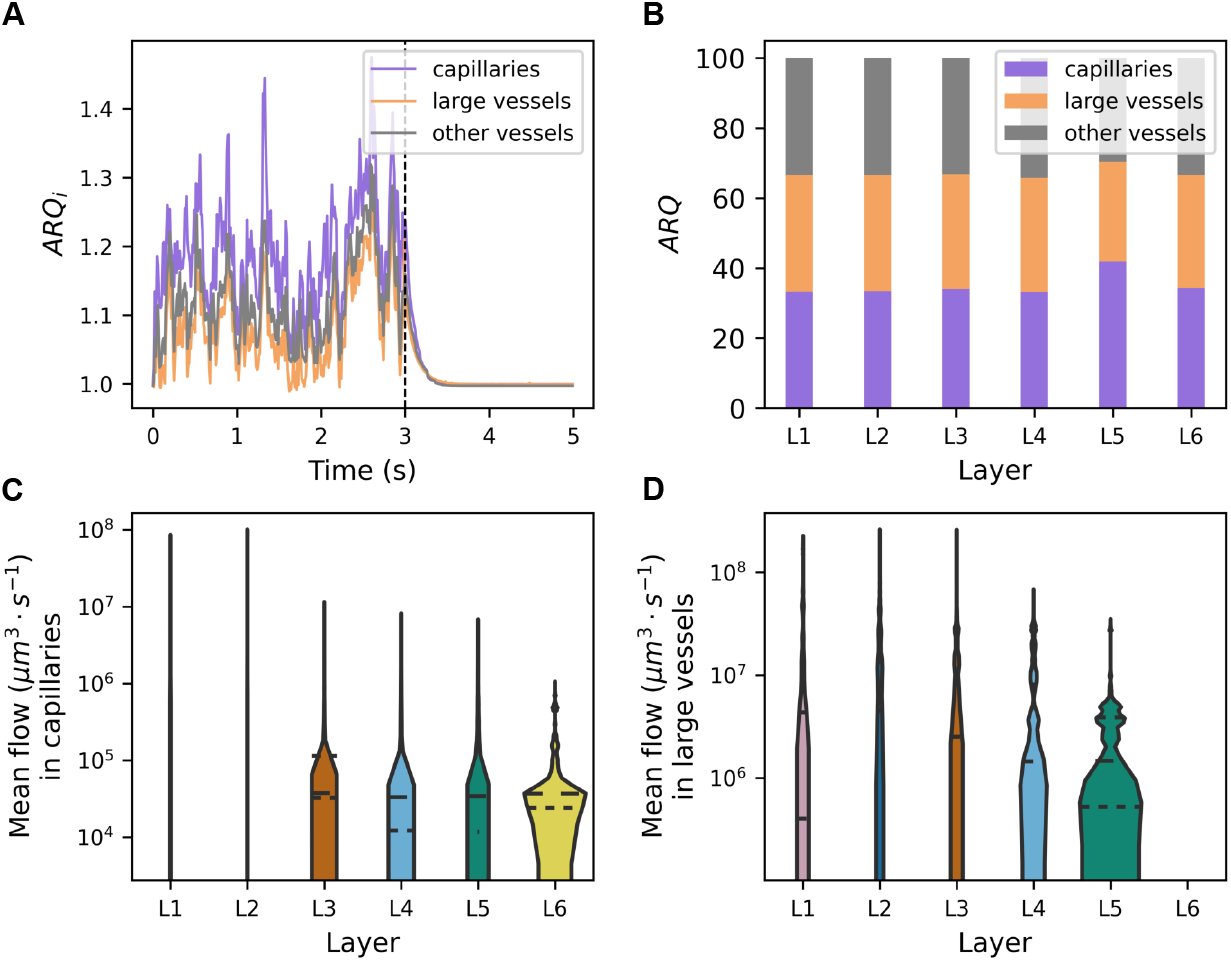
Layer-specific and localized astrocytic activity. (A) Average resting state flow ratio *ARQ*_*i*_, Eq (27), during a five-second period, with astrocytic stimulation occurring in the first three seconds. Average taken over large vessels is illustrated in orange, capillaries in purple, and other vessels in gray. (B) Percentage contribution of average resting state flow ratio *ARQ*, Eq (28), attributed to each vessel type averaged over the initial three seconds of stimulation. Each bar corresponds to a cortical layer, with orange representing variation due to large vessels, purple for capillaries, and gray for other vessels. (C) Violin plots presenting the distribution of mean flow over time across the six cortical layers in capillaries and in large vessels (D). Each violin corresponds to a specific cortical layer, and the height of the plot reflects the range of blood flow values within that layer. The width represents the number of segments in that range

Figure 5C-D present violin plots of the distribution of mean flow over time across the six cortical layers in capillaries (C) and in large vessels (D). Each violin corresponds to a specific cortical layer. The height of the plot reflects the range of blood flow values within that layer, while the width the number of segments in that range. Layer-specific values were more dispersed in L1 and L2. This may be due to the lower density of capillaries in L1 and L2, coupled with a higher prevalence of larger vessels. While astrocytic activity primarily affects capillaries, larger vessels such as arterioles and venules are still subject to astrocytic modulation [15,73,74], albeit to a lesser extent. The impact of astrocytes was represented by the maximum radius deformation. The simulation values were taken from Table 2 and Table 3. The larger diameter and distinct regulatory mechanisms of these vessels attenuate, but do not negate, the impact of astrocytic activity. Capillaries, in particular, are highly responsive to astrocytic signaling, with changes in astrocytic activity directly influencing their diameter and thus impacting blood flow dynamics [5].

### Impact of astrocytic activity on blood flow is spatially localized

To explore the effect of astrocytic activity on the local vasculature we looked at the flow variation induced by astrocytes in the neighboring segments. In Figure 6 we investigated the spatial range in which a given endfoot has an effect. To that end we computed the ratio of resting flow variation between a segment containing an endfeet and its neighbors of increasing order. We defined the segment order number *m* from an endfoot segment *k* the nth segments from both directions (see Section 2.8.1). The order ratio defined in Eq (29) is illustrated in Figure 6A by a gray dotted line. The ratio decreased exponentially with the order, indicating that the further a segment was from the endfeet, the less impact the endfeet activity had on the segment flow variation. Section 4 analyzed this effect further. We then chose a 0.1 threshold, below which the average order flow ratio was very low and considered negligible. We then introduced an upper bound limit *m* = 20, focusing only on the edges connected to endfeet up to *m* = 20 (see Figure 6A). For each edge in the vasculature, we perform the following steps:

**Figure 6.**
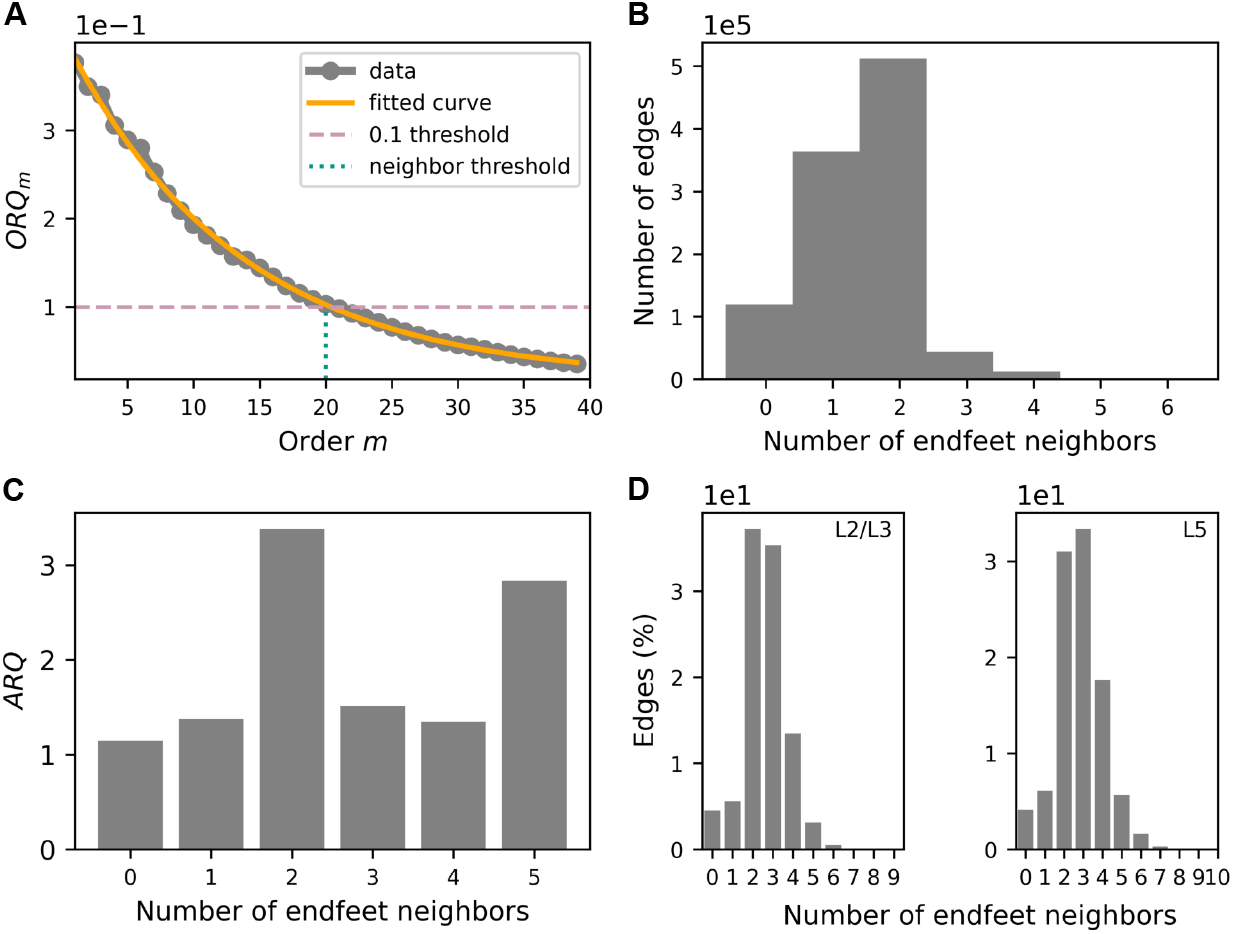
Effect of astrocytic activity on local vasculature. (A) Order flow ratio *ORQ*_*m*_, Eq (29), of the change in flow between edges linked to an endfoot and their adjacent edges (see Section 2.8). The x-axis shows the order *m*, defined as the number of segments starting from the one linked to an endfoot. Data is shown in gray (dotted line), a 0.1 threshold is indicated in pink (dash line), and neighbor threshold (order *m* = 20) is depicted in blue green (dotted line). The data can be fitted accurately by an exponential function (orange line): *f* (*m*) = *a e*^−*b m*^ + *c*. The final fitting parameters are *a* = 0.39, *b* = 0.07, and *c* = 0.01. (B) Distribution of the number of edges based on their proximity to the endfeet. The x-axis indicates the number of endfeet neighbors (close to each edge), while the y-axis represents the number of edges. (C) Variation in flow concerning the nearest endfeet. The x-axis represents the number of closest endfeet, while the y-axis illustrates the average resting state flow ratio *ARQ* defined in Eq (28). (D) Distribution of absolute flow variation with respect to the nearest endfeet. The x-axis denotes the number of nearest endfeet related to each edge above a 0.1 threshold (see A), while the percentage of edges is depicted on the y-axis. The left panel illustrates the distribution of flow across L2/L3, while the right panel displays the distribution of flow across L5.

- take an edge *k* in the vasculature
- consider all neighbors up to order 20 (see Section 2.8.1)
- count the number of endfeet in the neighbors
- classify the edge *k* into five categories depending on the number of endfeet (from zero to five, which is the maximum value we found) (see Figure 6B)
- consider the resting state ratio for the flow at each time for the edge *k* and take the mean.
- consider all edges *k* that belong to the same category and take the mean (see mean flow variation due to astrocytic activity (Figure 6C).)

We observed that segments from the vasculature were more likely to have one or two effective neighboring endfeet. Surprisingly, we found that the highest flow variation occurs when segments have two neighboring endfeet, rather than more (with the exception of the group of five neighboring endfeet). This unexpected result suggests that when astrocytes had more than two neighboring endfeet, their effects might sometimes cancel each other out, leading to less impact on flow variation (see Figure 6C). The high flow variation value for the group with five neighboring endfeet may be due in part to the relatively low number of segments in that group. Finally, we examined the distribution of the seven groups of segments in layers L2 and L3 in one hand and L5 (Figure 6D) in the other hand. The majority of edges in both L2/L3 and L5 had between two and three endfeet neighbors. In other words, the radii of the majority of edges are influenced by either two or three endfeet. This was fortuitous, as this two neighboring endfeet configuration coincides with the most effective configuration in terms of astrocytic control on blood flow variation (Figure 6C). Although the two neighboring endfeet configuration was slightly more likely in subgranular layers while the three neighboring endfeet configuration was slightly more likely in supragranular layers. Both configurations exhibited similar characteristics across all layers.

## 4. Discussion

We characterized a crucial mechanism of blood flow regulation in the brain: vasodilation of blood vessels due to astrocytic stimulation. We chose rat SSCx as the foundation of our studies due to its extensive coverage in the literature. The model consisted of several elements:

1. a piece of reconstructed vascular network from the rat SSCx that spanned several cortical layers (L1-L5 completely and L6 partially, [18]);
2. astrocytes carefully placed to populate the space surrounding the vasculature as well as their endfeet to complete the coupling with the vascular network [8]; and
3. a set of differential equations (see [**?**]) coupling the astrocyte activity (i.e. the radius dilation caused by the endfeet) and the resulting variation in blood flow values.

We selected a ROU process to model the dynamics of vessel radius changes due to astrocytic activity. To accurately characterize flow variations in response to radius changes, we defined new metrics: the resting state ratios. These metrics were applied to both radius and blood flow (See 2). Experimental data on radius ratios for vessel segments exclusively linked to endfeet is sparse [14,39,41–43,61–63] because measuring radius changes requires simultaneous measurements in several vessels. We use these literature data as constraints for our model to set the maximum vasodilation for capillaries (38%) and large vessels (23%). This constraint induced in our simulations to produce a near-unity flow ratio in most segments connected to endfeet, indicating a high degree of stability in flow rates even during the initial astrocytic activity period. The distribution of radius ratios in Figure 3A follows a smooth trend, indicating that vessel diameter changes occur predictably over time. In contrast, the distribution of flow ratios in Figure 3B appears scattered, highlighting a highly non linear relationship between vessel radius changes and resulting flow variations. The evolution of radius and flow averaged over all vessel segments revealed transient increases during stimulation, followed by a return to baseline levels in the passive phase (see Figure 3D-F). The vasodilation-induced mean flow increase is possibly mediated by astrocytes, consistent with their established role in regulating cerebral blood flow. Together, these results imply that astrocytes help stabilize flow dynamics, ensuring consistent cerebral blood flow. In turn, this ensures a reliable supply of oxygen and nutrients to neurons [64,75], prevents ischemic conditions, and supports metabolic homeostasis [75,87,88].

We validated our model simulations by addressing two key aspects:

1. the comparison of flow values in representative vessels (capillaries and large vessels, Figure 4A) to biological measurements, and,
2. the response of flow to time-dependent vasodilation induced by astrocytic activity (Figure 4C-E).

Quantifying flow in a dense, interconnected microvascular network like the rat SSCx is experimentally challenging [44]. While our simulations produced higher flow values than reported experimentally, several factors might explain this. First, simulations may capture physiological phenomena that are difficult to measure directly. Second, flow measurement techniques can introduce artifacts; for example, detection devices may consume a small portion of the flow, affecting accuracy. Third, the sample size differences are substantial: our model simulates numerous segments, whereas experimental studies typically analyze fewer than 50 segments.[67–69,89–93]. Lastly, our model does not account for the biphasic nature of flow due to red blood cells (RBCs). For computational simplicity, we assumed a fixed RBC volume [70] and hematocrit proportion [71], which may not capture the effects of RBC compression in capillaries, where smaller diameters can alter RBC volume and flow dynamics. Modeling this compression and incorporating vasoconstriction would likely yield a more symmetric and physiologically accurate flow distribution [94]. Indeed, our model only accounts for vasodilation and disregards vasoconstriction. If we modeled both vasodilation and vasoconstriction, we might have expected the flow distribution to be more symmetric around the resting state ratio. This is because vasoconstriction would balance the effects of vasodilation, potentially smoothing out the extremes and leading to a more even distribution of flow values. The distribution of flow and velocity values might show additional peaks corresponding to the points where vasoconstriction counteracts vasodilation (see Figure 4C-F). However, this does not necessarily mean that it would be smoother; rather, it would be more balanced, reflecting the bidirectional modulation of vessel radii. The overall shape could become more complex, indicating the interplay between constrictive and dilative forces on flow dynamics. Schmid et al. demonstrated the impact of RBCs on the microvascular perfusion and highlighted the heterogeneity of changes in RBCs distribution [94].

It is crucial to acknowledge these potential sources of discrepancy when interpreting the disparities between simulated and literature values. By comparing the simulation results (see Figure 4A and Figure 4B) with the literature data [14–17,95] we showed that the mean values of simulated flow and velocity closely aligned with the reported values in the literature, with the majority of data falling within the typical range observed for rats. Indeed, the ranges from the model include the ranges in the literature and can be reproduced by the model.

Simulations showed that the highest variability of blood flow (67%) happens inside capillaries. This underscores the pivotal role of capillaries in fine-tuning cerebral microvasculature flow dynamics [14]. Additionally, this aligns with the widely accepted notion that capillaries are key sites for modulating blood flow to meet the metabolic demands of neural tissue. The morphology of the vasculature led to distinct patterns of layer-specific flow distribution. Deeper layers (L4-L5-L6), with higher capillary contribution, exhibited narrower flow value ranges, indicating tighter regulation [32,36,94,96]. Conversely, upper layers, abundant in larger vessels, showed greater flow fluctuations, likely influenced by capillary dilation [25,97–99].

We introduced the *ORQ*_*m*_ metric (Eq (29)) to quantify astrocytic activty effects on the local vasculature. We observe a reduction of this ratio with increasing segment order (distance). It may be attributed to the dispersion of the perturbation caused by the branching of the segments (see Figure 6A). This analysis showed that astrocytes have localized effects on nearby blood vessels (i.e. on the 20 first neighbors considering an arbitrary threshold of 10% Figure 6A).

Despite their widespread presence, a comprehensive understanding of endfoot physiology is still lacking. This gap in knowledge is partly due to the limited identification and characterization of the specific proteins that differentiate the endfoot from the rest of the astrocyte body. Our findings suggested a relationship between astrocytic coverage and blood flow regulation where moderate astrocytic presence (two or three effective endfeet) elicits a robust response, while excessive coverage (more than three endfeet) leads to diminished effects (see Figure 6C-D). This observation is supported by the literature [100]. More precisely, the interaction between multiple neighboring endfeet may present diminishing returns. Instead, the collective influence of multiple neighboring endfeet can lead to complex interactions, constructive or destructive interference, and heightened flow variability in certain scenarios. Finally, if endfeet are spatially clustered or arranged in a specific pattern, such setup can contribute to higher flow variability. Layer-specific configurations influencing flow regulation highlight the intricate interplay between cellular components and vascular function across cortical regions. Layer-specific endfoot configurations further modulated flow, with supragranular layers optimized for two endfeet and subgranular layers for three [101,102].

Although it is assumed in our model that capillary diameters can change dynamically with activity, it is important to note that this remains a subject of debate in the literature. Evidence suggests that the first vessels off a penetrating arteriole may exhibit rapid dilation [14], while capillaries in the middle of the network tend to show slower or negligible dilation on the scale of seconds and only respond significantly over longer periods, such as hundreds of seconds [80,81].

Our current modeling approach focuses solely on astrocytic mechanisms and the effect of endfeet on vessel diameters. It relies on simplifying assumptions and parameters derived from experimental data. Therefore, there are still many avenues for improvement yet to be explored. We acknowledge that the neurovascular unit is highly complex and includes additional mechanisms of blood flow regulation that our model does not explicitly address. For instance, recent studies have shown that pericytes, positioned at capillary junctions, can dynamically modulate blood flow through contractile activity and signaling mechanisms that propagate along the endothelium ([76]). These findings highlight the importance of pericyte-mediated hemodynamic control, which our model does not incorporate. Furthermore, ion channel activity in pericytes, including inward-rectifier *K*^+^ channels and voltage-dependent channels, has been shown to modulate capillary hemodynamics, further contributing to cerebral blood flow regulation [77]. Despite sharing molecular machinery with smooth muscle cells, pericytes exhibit distinctive ion channel properties that influence cerebral blood flow differently. Although these mechanisms are integral to the neurovascular unit’s functioning, our model does not explicitly incorporate pericyte dynamics, as the focus remains on astrocyte-mediated regulation of vessel diameters. Future iterations of our model should aim to incorporate upward signal transfer within the endothelium, as well as the integration of pericyte and endothelial signaling to better represent the neurovascular unit’s network effects and to simulate more realistic flow dynamics. Incorporating these network effects would allow for a more detailed analysis of how local changes propagate and influence the overall flow distribution across cortical layers. By acknowledging these limitations and areas for improvement, our study lays the groundwork for more sophisticated models that include pericyte dynamics, endothelial signaling, and bidirectional vessel modulation, which would provide a more comprehensive understanding of neurovascular coupling.

Despite the alignment with anatomical observations, our approach represents a simplification compared to more advanced hemodynamic models that account for distributed inflows across finer anatomical details. Such models could improve the physiological realism but would require significantly more computational resources and detailed parameterization. Future research could explore incorporating these features, such as shear-dependent flow distribution and spatially distributed inflows, to enhance the fidelity of the model.

The model predictions could be validated experimentally using *in vivo* imaging techniques and manipulations of astrocytic activity. Furthermore, the model could be expanded to the full mouse brain vasculature dataset by Ji et al. [103]. This would require repairing the vasculature gap and label the different types of vessels based on the Xiong et al. atlas [104]. Finally, extending the model to pathological conditions associated with altered neurovascular coupling, such as neurodegenerative diseases and stroke, could provide insights into disease mechanisms and potential therapeutic targets.

## 5. Conclusions

In conclusion, the study sheds light on the intricate interplay between astrocytic signaling and cerebral blood flow dynamics. We provide a useful framework for future research into physiological and pathological conditions associated with neurovascular dysfunctions.

## Author Contributions

Conceptualization, S.B., N.C., and D.K.; methodology, S.B. and N.C.; software, S.B., N.C., C.K. and A.A.; validation, S.B. and N.C.; formal analysis, S.B.; investigation, S.B., N.C., C.K., Y.R.; resources, H.M. and D.K.; data curation, S.B.; writing—original draft preparation, S.B., N.C., C.K., Y.R., A.C., A.A., C.F., S.A., H.M., and D.K.; writing—review and editing, S.B., N.C., C.K., Y.R., A.C., A.A., C.F., S.A., H.M., and D.K.; visualization, C.F.; supervision, S.B. and D.K.; project administration, S.B., D.K. and H.M.; funding acquisition, H.M. All authors have read and agreed to the published version of the manuscript

## Funding

This study was funded by the Blue Brain Project, a research center of the École polytechnique fédérale de Lausanne (EPFL), from the Swiss Government’s ETH Board of the Swiss Federal Institutes of Technology.

## Acknowledgments

Bruno Weber is gratefully acknowledged for providing the rat SSCx network. We would like to thank Thomas Delemontex, Eleftherios Zisis, and Tristan Carel for their invaluable contributions in providing engineering support. We extend our profound thanks to all members of the molecular systems team for their insightful comments and invaluable discussions. Additionally, we are grateful to Dimitri Rodarie for his constructive feedback and engaging discussion. We finally thank Jean-Denis Courcol and Claudia Savoia for organizing web portal development and James Gonzalo King for the HPC support.

## Appendix A

### Appendix A.1 Exit flow computation

Our current approach for calculating flow at the exit nodes (presented in Section 2.3) is a naive approach that simply distributes the total entry flow among the exit nodes proportionally to their respective area.

Here we present an alternative approach that we considered at the beginning of our analysis, but which we later decided to discard because it required too many computational resources. In this section we consider weights proportional to the effective conductance *c*_*ab*_ between an entry node *a* and an exit node *b*

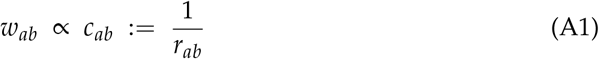

The effective conductance is the inverse of the effective resistance. Incidentally, this is a popular distance measure used in graph theory and electrical networks [**?**].

If we associate each node *i* with an element *e*_*i*_ of the standard basis of the Euclidean vector space ℝ^*n*^, we can define the effective resistance between any nodes *i, j* as follows:

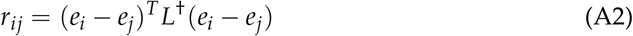

where *L*^†^ is the pseudo-inverse of the graph Laplacian matrix. The Laplacian matrix is a singular matrix, and therefore we need to consider the pseudo-inverse.

The weight definition proposed in this section is more appropriate as it considers the overall effective conductance between an entry node and an exit node, which depends on the length and radius of each edge within the graph, rather than solely relying on the area of the exit node.

Our graph contains 10^6^ nodes, out of which 19004 are boundary nodes. As we have chosen three entry nodes, the remaining 19001 are exit nodes. Given these values, even when employing optimized parallel algorithms on distributed systems, explicitly calculating the pseudoinverse of the Laplacian matrix becomes exceedingly time consuming.

Consequently, we have opted to compute solely the effective resistance between the entry node *a* and the exit node *b*, ignoring the values of *r*_*ij*_ between nodes that we are not interested in. If we call *v*_*ab*_ := *e*_*a*_− *e*_*b*_, we can efficiently solve the linear system *Lx* = *v*_*ab*_ to obtain 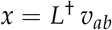, and then obtain 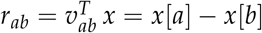.

Although this approach is considerably faster than calculating the pseudoinverse, it remains highly time-consuming as it necessitates the solution of 3 × 19001 linear systems, and cannot be used in practice. Due to this constraint, we have opted for the simpler area-based approach, which nonetheless, yields comparable results.

As depicted in A1, we observe that effective resistance exhibits a decreasing relationship with area. Although this functional relationship is not linear, indicating that the two approaches yield very different weights, this coarse analysis provides evidence that greater area corresponds to an increased effective conductance. Thus, we employ this heuristic argument to justify our chosen approach.

**Figure A1.**
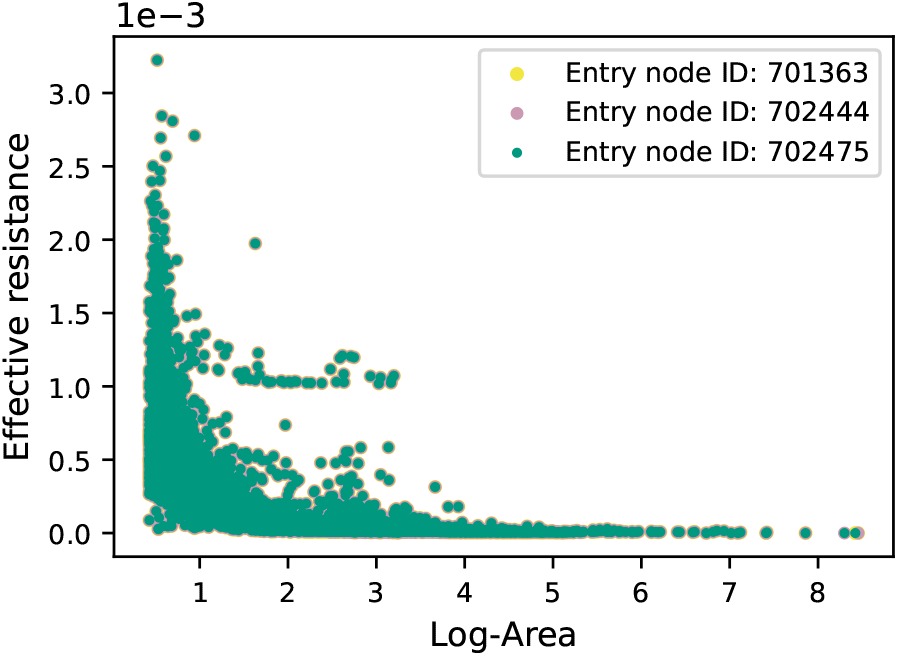
Effective resistance. A scatter plot depicting the effective resistance (ER) as a function of the logarithm of the exit node area. This analysis involves the calculation of effective resistance between each of the three selected entry nodes and all exit nodes in the network.

## Appendix B Model validation

Validation focuses solely on the blood vessels in the internal part of the vasculature, within the defined bounds of [*x*_*min*_, *x*_*max*_], [*y*_*min*_, *y*_*max*_], and [*z*_*min*_, *z*_*max*_] with *x*_*min*_ = 67.70 *µm, x*_*max*_ = 622.30 *µm, y*_*min*_ = 1033.50 *µm, y*_*max*_ = 1684.99 *µm, z*_*min*_ = 273.27 *µm*, and *z*_*max*_ = 926.73 *µm*. So, approximately 17% of the vasculature falls within the defined region. In this way, we minimize the possible distortions introduced by the boundary conditions and take into account contributions from all nodes in the vasculature.

## Appendix C Choice of time-step

In order to select an appropriate time step for our simulation we consider the time evolution of the mean radius in the vasculature, as shown in Figure **??**. The objective is to choose a time step that is not too large. We observe that the slope of the curve becomes nearly flat after time *t* ≈ 0.4. The region where the curve is almost flat is usually called the steady state, or plateau. Considering a time step size of Δ*t* = 0.01, we estimate that approximately 40 time steps are required to reach the plateau (see Figure **??**).

The shape of the curve representing the average radius evolution during stimulation can be described by the following parametric function

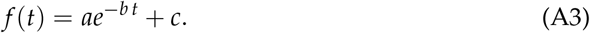

## Appendix D Passive properties of the network

The passive property of the network refers to the intrinsic characteristics of the vascular network. The intrinsic characteristics are responsible for the dynamics of the radii and flow inside the vasculature when astrocytic activity ends. This is what we call the passive activity (i.e. the passive response of the vasculature). In our model the phase corresponding to the passive response of the vasculature can be described by Equation **??** by setting *σ* = 0. By defining *τ* as the time at which stimulation ceases, after setting *σ* = 0 in Equation **??**, we get the subsequent dynamics:

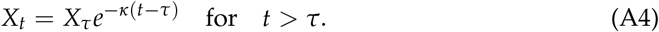

Here, *X*_*t*_ represents the blood vessel radius at time *t, X*_*τ*_ is the blood vessel radius at the cessation of stimulation *τ* and *κ* governs the rate of decay. This equation describes the exponential decay of the blood vessel radius after the cessation of astrocytic stimulation.

The average radius can be determined through

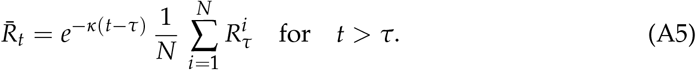

Here 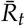 represents he average blood vessel radius at time *t* after the cessation of stimulation. The sum 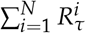 calculates the initial radii of *N* blood vessels *segment*_*v*_*essel*_*o*_*ver*_*t*_*imes* 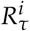 at the time of stimulation cessation. The exponential decay factor accounts for the decrease in blood vessel radius over time, reflecting the continued influence of astrocytic activity withdrawal. These formulas show how blood vessel radius evolves following the cessation of astrocytic stimulation.

**Figure A2.**
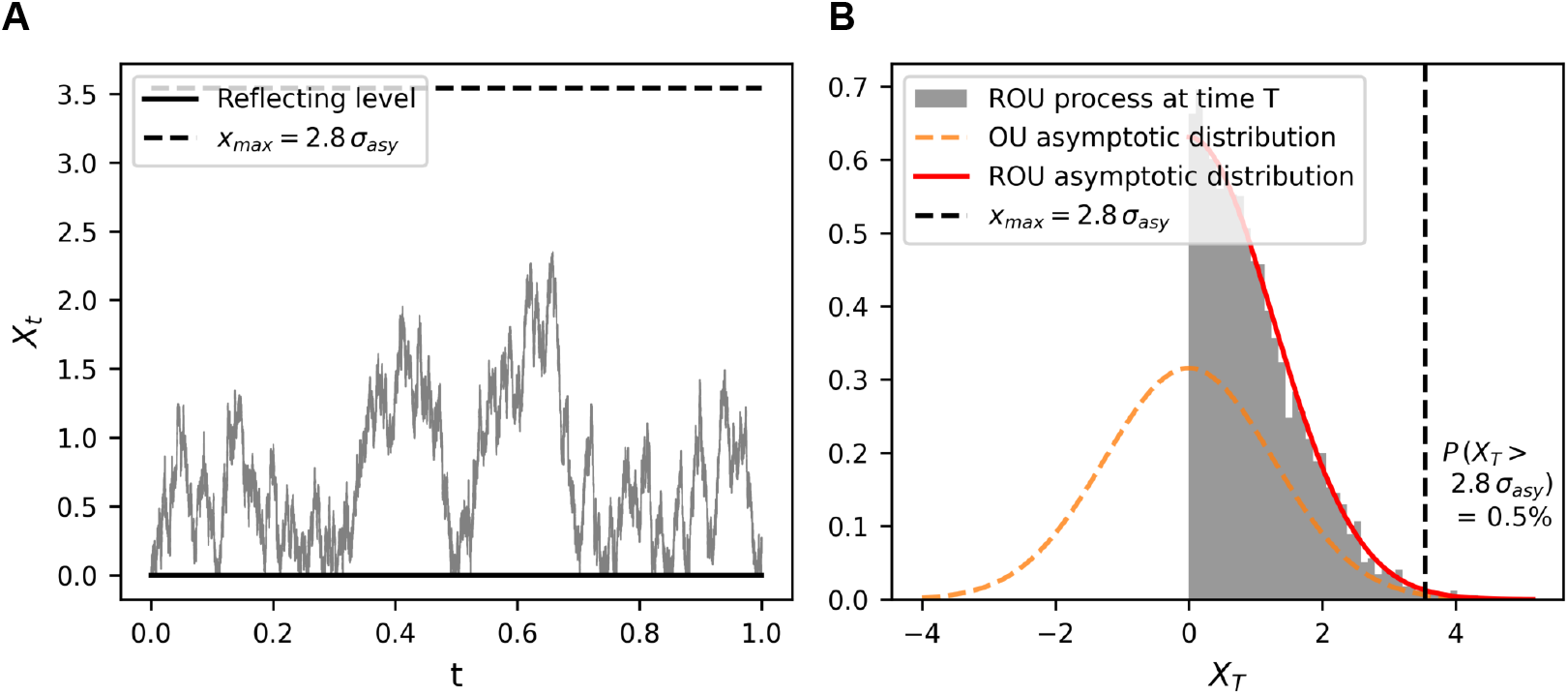
Calibration of the Ornstein-Uhlenbeck (OU) process. (A) Sample path of a Reflected OU (ROU) process with *T* = 1, *κ* = 5 and *σ* = 4. The solid black line represents the reflecting model, while the dashed black line denotes a realistic maximum value for the ROU process. (B) Comparison of the asymptotic OU and ROU distributions. We simulated 5,000 ROU paths and constructed a histogram using the values of *X*_*T*_ at *T* = 1. The solid red line represents the ROU asymptotic distribution, the dashed red line represents the OU asymptotic distribution, and the dashed black line denotes a realistic maximum value for the ROU process.

**Figure A3.**
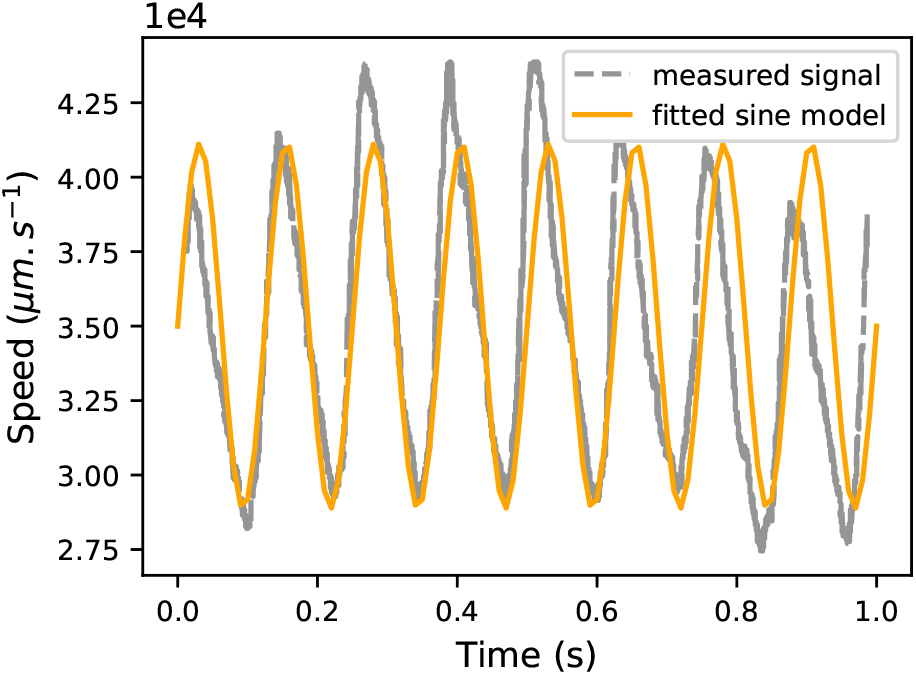
Fit of a sinusoidal blood flow signal. Comparison of blood velocity signal (gray dashed line) obtained from [**?**], with a fitted sine function *v*(*t*) = *A* sin(2 *π f t*) + *C* (orange continuous line) with *A* = 6119 *µm* · *s*^−1^, *f* = 8 *s*^−1^, and *C* = 35, 000 *µm* · *s*^−1^.

**Figure A4.**
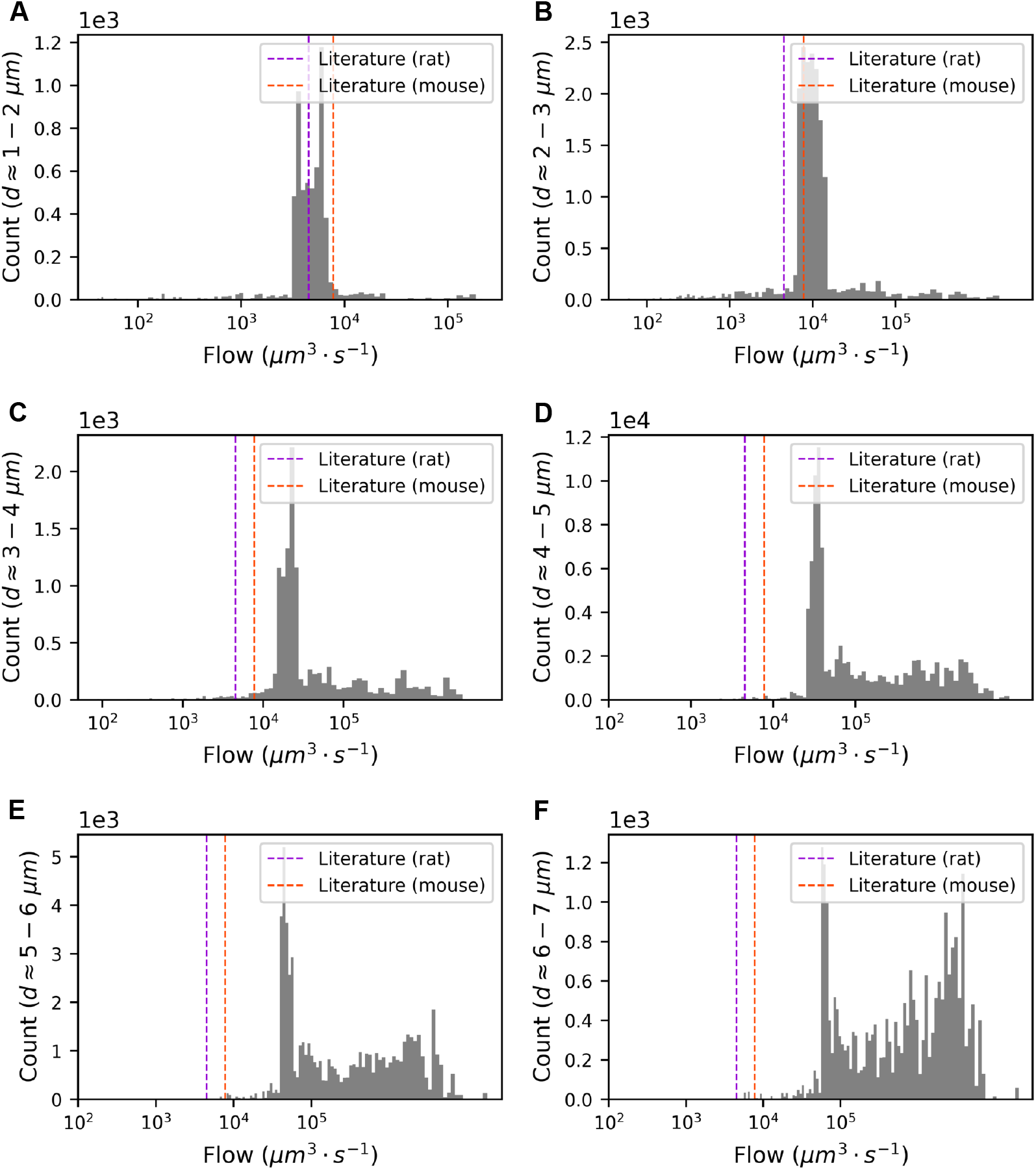
Validation of simulation results against literature data. Distribution of blood flow in capillaries categorized by diameter plotted on a logarithmic scale, with dashed lines from previous studies (see Section B). (A) 1-2 *µm*, (B) 2-3 *µm*, (C) 3-4 *µm*, (D) 4-5 *µm*, (E) 5-6 *µm*, (F) 6-7 *µm*.

**Figure A5.**
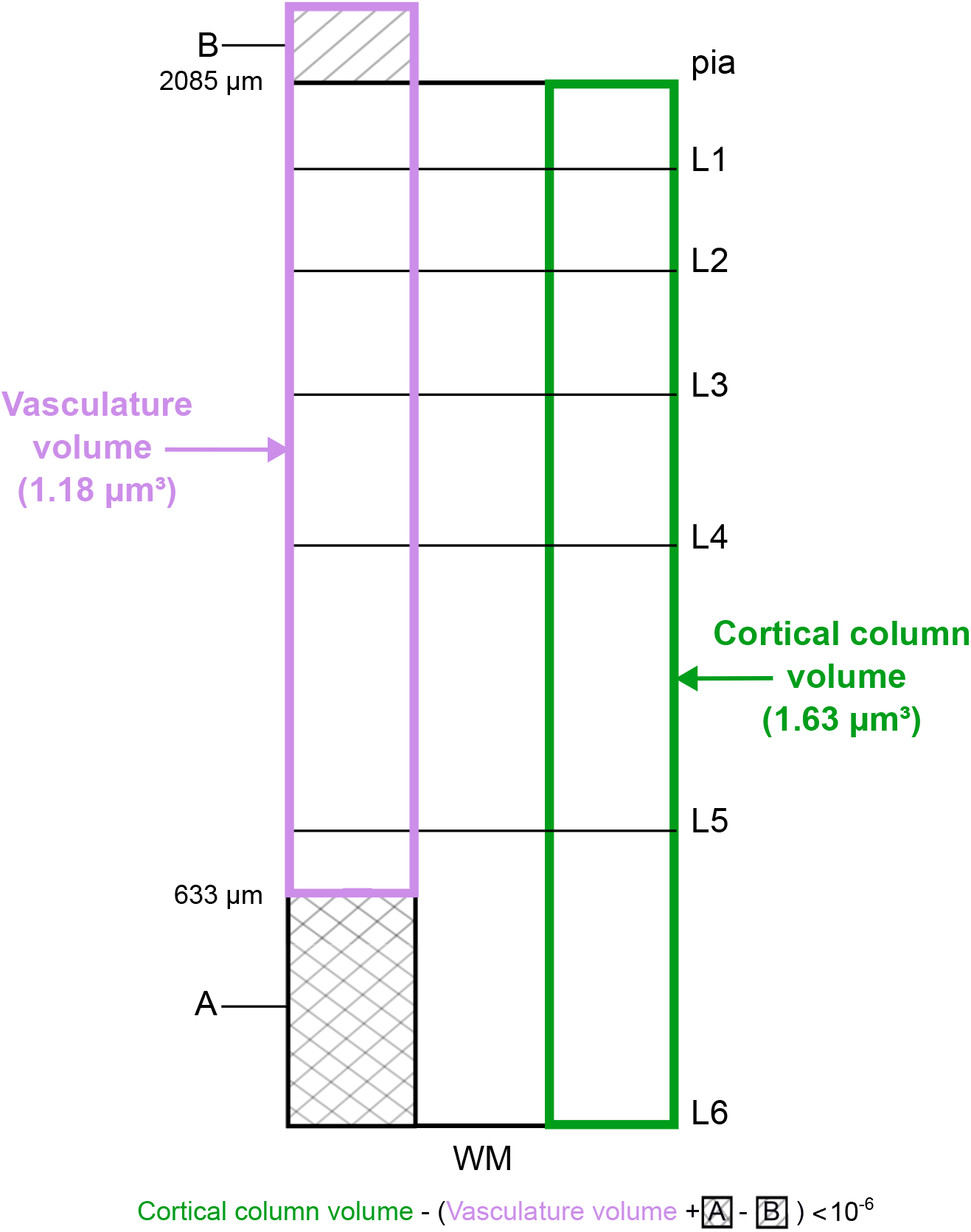
Volume discrepancy: misalignment between vasculature and microcircuit in the rat SSCx. The pink rectangle represents the vasculature volume, while the green one shows the cortical column volume. The B striped volume represents the volume of the vasculature above L1, while the A hatched volume represents the missing vasculature from three-fourths of L6 down to the white matter (WM).

**Figure A6.**
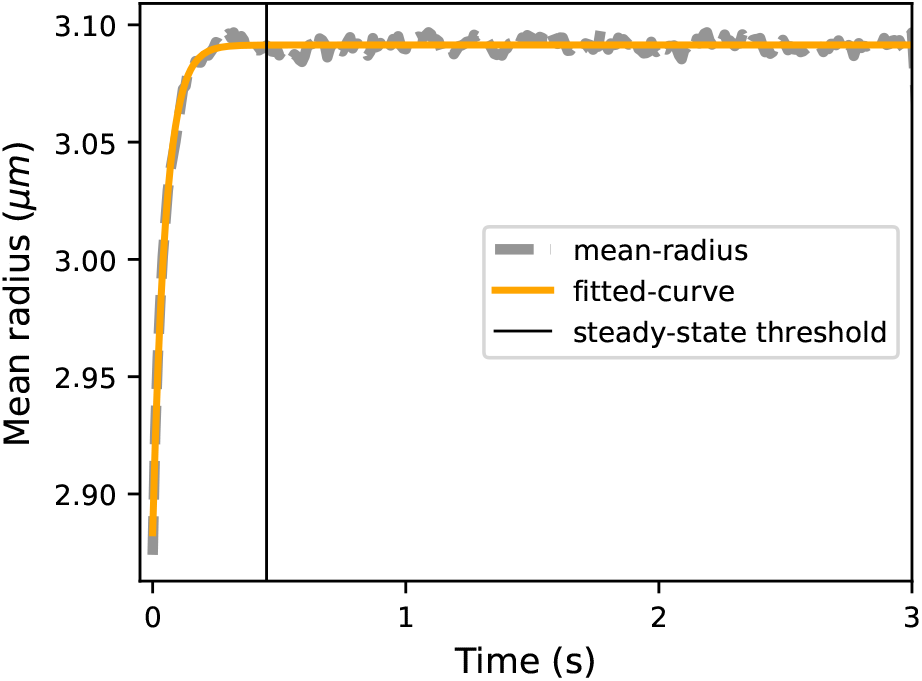
Dynamical evolution of the average radii. Comparison of the average radii over time (gray dashed line), with a fitted exponential function *f* (*t*) = *ae*^−*b t*^ + *c* (orange continuous line) with fitted parameters *a* = −0.23, *b* = 21.38, and *c* = 3.03.

**Table A1.** Validation of the simulation results against data from the literature. Units of flow values are expressed in *µm*^3^ · *s*^−1^ ·^*^ [69]

|  | Flow in capillaries |  | Flow in large vessels |  |
| --- | --- | --- | --- | --- |
|  | average | range | average | range |
| Literature | $6.65 \cdot 10^3$ | $4.85 \cdot 10^3 - 7.78 \cdot 10^3$ | - | $1 \cdot 10^5 - 1 \cdot 10^7$ * |
| Our network | $4.93 \cdot 10^5$ | $0.01 - 1.83 \cdot 10^7$ | $2.28 \cdot 10^7$ | $1.54 \cdot 10^5 - 1.59 \cdot 10^8$ |

**Table A2.**
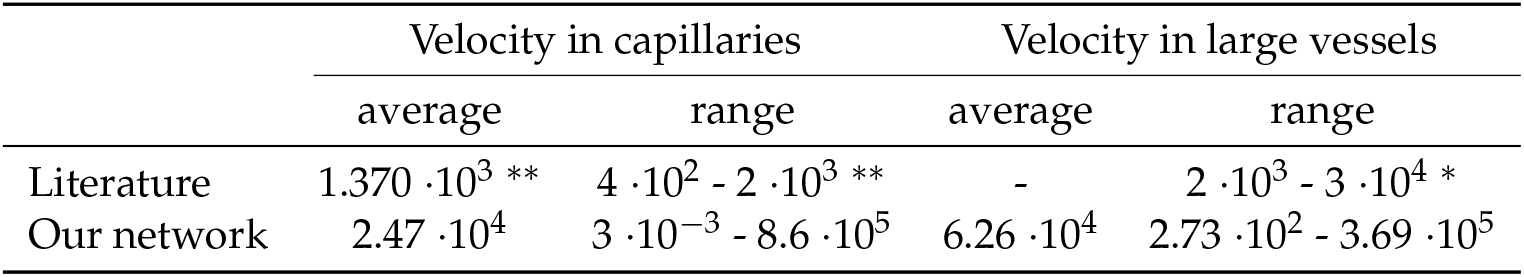
Validation of the simulation results against data from the literature. Units of velocity values are expressed in *µm* · *s*^−1. **^ [67,68,89–93], ^*^[69]

|  | Velocity in capillaries |  | Velocity in large vessels |  |
| --- | --- | --- | --- | --- |
|  | average | range | average | range |
| Literature | $1.370 \cdot 10^3$ ** | $4 \cdot 10^2 - 2 \cdot 10^3$ ** | - | $2 \cdot 10^3 - 3 \cdot 10^4$ * |
| Our network | $2.47 \cdot 10^4$ | $3 \cdot 10^{-3} - 8.6 \cdot 10^5$ | $6.26 \cdot 10^4$ | $2.73 \cdot 10^2 - 3.69 \cdot 10^5$ |

**Table A3.** Unit conversions and parameters for blood flow calculations.

| Parameter | Value / Conversion |
| --- | --- |
| Flow rate in large vessels | $\text{nl} \cdot \text{s}^{-1}$ |
| Flow rate in capillaries | $\text{RBC} \cdot \text{s}^{-1}$ |
| Velocity | $\text{mm} \cdot \text{s}^{-1}$ |
| Millimeter to micrometer | $1 \text{ mm} = 10^3 \mu\text{m}$ |
| Femtoliter to cubic micrometer | $1 \text{ fL} = 1 \mu\text{m}^3$ |
| RBC volume conversion | $\frac{\text{RBCv}}{\text{hematocrit}} \cdot \text{RBC}$ |
| RBC volume (RBCv) | $56.51 \text{ fL}$ [70] |
| Hematocrit proportion | 0.45 [71] |

where RBCv indicates the volume of a single red blood cell. Baskurt et al. [70] reports a RBCv of 56.51 *f l* in rats. According to Pries et al. [71] the hematocrit proportion is 0.45.

## Disclaimer/Publisher’s Note

The statements, opinions and data contained in all publications are solely those of the individual author(s) and contributor(s) and not of MDPI and/or the editor(s). MDPI and/or the editor(s) disclaim responsibility for any injury to people or property resulting from any ideas, methods, instructions or products referred to in the content.

## References

1. Verkhratsky, A.; Toescu, E. Neuronal-glial networks as substrate for CNS integration. J. Cell Mol. Med. 2006, 10, 869–879.

2. Magistretti, P.; Allaman, I. Lactate in the brain: from metabolic end-product to signalling molecule. Nat. Rev. Neurosci. 2018, 19, 235–249.

3. Kacem, K.; Lacombe, P.; Seylaz, J.; Bonvento, G. Structural organization of the perivascular astrocyte endfeet and their relationship with the endothelial glucose transporter: A confocal microscopy study. Glia 1998, 23, 1–10.

4. Abbott, N.; Rönnbäck, L.; Hansson, E. Astrocyte–endothelial interactions at the blood–brain barrier. Nat. Rev. Neurosci. 2006, 7, 41–54.

5. Mathiisen, T.; Lehre, K.; Danbolt, N.; Ottersen, O. The perivascular astroglial sheath provides a complete covering of the brain microvessels: An electron microscopic 3D reconstruction. Glia 2010, 58, 1094–1103.

6. Simard, M.; Arcuino, G.; Takano, T.; Liu, Q.; Nedergaard, M. Signaling at the gliovascular interface. J. Neurosci. 2003, 23, 9254–9262.

7. Korogod, N.; Petersen, C.C.; Knott, G.W. Ultrastructural analysis of adult mouse neocortex comparing aldehyde perfusion with cryo fixation. eLife 2015, 4, e05793.

8. Zisis, E.; Keller, D.; Kanari, L.; Arnaudon, A.; Gevaert, M.; Delemontex, T.; et al. Digital Reconstruction of the Neuro-Glia-Vascular Architecture. Cereb. Cortex 2021, 1, 1–18.

9. Kenny, A.; Plank, M.; David, T. The role of astrocytic calcium and TRPV4 channels in neurovascular coupling. J. Comput. Neurosci. 2018, 44, 97–114.

10. Wieronska, J.; Cieślik, P.; Kalinowski, L. Nitric Oxide-Dependent Pathways as Critical Factors in the Consequences and Recovery after Brain Ischemic Hypoxia. Biomolecules 2021, 11, 1097.

11. Schiera, G.; Di Liegro, C.; Schirò, G.; Sorbello, G.; Di Liegro, I. Involvement of Astrocytes in the Formation, Maintenance, and Function of the Blood-Brain Barrier. Cells 2024, 13, 150.

12. Mester, J.; Rozak, M.; Dorr, A.; Goubran, M.; Sled, J.; Stefanovic, B. Network response of brain microvasculature to neuronal stimulation. Neuroimage 2024, 287, 120512.

13. Djurich, S.; Secomb, T. Analysis of potassium ion diffusion from neurons to capillaries: Effects of astrocyte endfeet geometry. Eur. J. Neurosci. 2024, 59, 323–332.

14. Hall, C.; Reynell, C.; Gesslein, B.; Hamilton, N.; Mishra, A.; Sutherland, B.; et al. Capillary pericytes regulate cerebral blood flow in health and disease. Nature 2014, 508, 55–60.

15. Mishra, A.; Reynolds, J.; Chen, Y.; Gourine, A.; Rusakov, D.; Attwell, D. Astrocytes mediate neurovascular signaling to capillary pericytes but not to arterioles. Nat. Neurosci. 2016, 19, 1619–1627.

16. Kisler, K.; Nelson, A.; Rege, S.; Ramanathan, A.; Wang, Y.; Ahuja, A.; et al. Pericyte degeneration leads to neurovascular uncoupling and limits oxygen supply to brain. Nat. Neurosci. 2017, 20, 406–416.

17. Rungta, R.; Chaigneau, E.; Osmanski, B.; Charpak, S. Vascular compartmentalization of functional hyperemia from the synapse to the pia. Neuron 2018, 99, 362–375.e4.

18. Reichold, J.; Stampanoni, M.; Keller, L.; Buck, A.; Jenny, P.; Weber, B. Vascular graph model to simulate the cerebral blood flow in realistic vascular networks. J. Cereb. Blood Flow Metab. 2009, 29, 1429–1443.

19. Payne, S.; El-Bouri, W. Modelling dynamic changes in blood flow and volume in the cerebral vasculature. NeuroImage 2018, 176, 124–137.

20. Gagnon, L.; Sakadzic, S.; Lesage, F.; Musacchia, J.; Lefebvre, J.; Fang, Q.; et al. Quantifying the microvascular origin of BOLD-fMRI from first principles with two-photon microscopy and an oxygen-sensitive nanoprobe. J. Neurosci. 2015, 35, 3663–3675.

21. Gagnon, L.; Smith, A.; Boas, D.; Devor, A.; Secomb, T.; Sakadzic, S. Modeling of cerebral oxygen transport based on in vivo microscopic imaging of microvascular network structure, blood flow, and oxygenation. Front. Comput. Neurosci. 2016, 10, 82.

22. Lorthois, S.; Cassot, F.; Lauwers, F. Simulation study of brain blood flow regulation by intra-cortical arterioles in an anatomically accurate large human vascular network; Part I: Methodology and baseline flow. NeuroImage 2011, 54, 1031–1042.

23. Boas, D.; Jones, S.; Devor, A.; Huppert, T.; Dale, A. A vascular anatomical network model of the spatio-temporal response to brain activation. NeuroImage 2008, 40, 1116–1129.

24. Safaeian, N.; Sellier, M.; David, T. A computational model of hemodynamic parameters in cortical capillary networks. J. Theor. Biol. 2011, 271, 145–156.

25. Gould, I.; Tsai, P.; Kleinfeld, D.; Linninger, A. The capillary bed offers the largest hemodynamic resistance to the cortical blood supply. J. Cereb. Blood Flow Metab. 2017, 37, 52–68.

26. Hartung, G.; Badr, S.; Mihelic, S.; Dunn, A.; Cheng, X.; Kura, S.; et al. Mathematical synthesis of the cortical circulation for the whole mouse brain-part II: Microcirculatory closure. Microcirculation 2021, 28, e12687.

27. Linninger, A.; Hartung, G.; Badr, S.; Morley, R. Mathematical synthesis of the cortical circulation for the whole mouse brain-part I: Theory and image integration. Comput. Biol. Med. 2019, 110, 265–275.

28. Kopylova, V.; Boronovskiy, S.; Nartsissov, Y. Fundamental constraints of vessel network architecture properties revealed by reconstruction of a rat brain vasculature. Math. Biosci. 2019, 315, 108237.

29. Roy, T.; Secomb, T. Functional implications of microvascular heterogeneity for oxygen uptake and utilization. Physiol. Rep. 2022, 10, e15303.

30. Attwell, D.; Buchan, A.; Charpak, S.; Lauritzen, M.; MacVicar, B.; Newman, E. Glial and neuronal control of brain blood flow. Nature 2010, 468, 232–243.

31. Howarth, C. The contribution of astrocytes to the regulation of cerebral blood flow. Front. Neurosci. 2014, 8, 103.

32. MacVicar, B.; Newman, E. Astrocyte regulation of blood flow in the brain. Cold Spring Harb. Perspect. Biol. 2015, 7, a020388.

33. Howarth, C.; Sutherland, B.; Choi, H.; Martin, C.; Lind, B.; Khennouf, L.; et al. A critical role for astrocytes in hypercapnic vasodilation in brain. J. Neurosci. 2017, 37, 2403–2414.

34. Tesler, F.; Linne, M.; Destexhe, A. Modeling the relationship between neuronal activity and the BOLD signal: Contributions from astrocyte calcium dynamics. Sci. Rep. 2023, 13, 6451.

35. Brazhe, A.; Verisokin, A.; Verveyko, D.; Postnov, D. Astrocytes: New evidence, new models, new roles. Biophys. Rev. 2023, 15, 1303–1333.

36. Cohen, Z.; Molinatti, G.; Hamel, E. Astroglial and vascular interactions of noradrenaline terminals in the rat cerebral cortex. J. Cereb. Blood Flow Metab. 1997, 17, 894–904.

37. Hösli, L.; Zuend, M.; Bredell, G.; Zanker, H.; Porto de Oliveira, C.; Saab, A.; et al. Direct vascular contact is a hallmark of cerebral astrocytes. Cell Rep. 2022, 39, 110599.

38. Takahashi, S. Metabolic contribution and cerebral blood flow regulation by astrocytes in the neurovascular unit. Cells 2022, 11, 813.

39. Devor, A.; Tian, P.; Nishimura, N.; Teng, I.; Hillman, E.M.C.; Narayanan, S.N.; et al. Suppressed neuronal activity and concurrent arteriolar vasoconstriction may explain negative blood oxygenation level-dependent signal. J. Neurosci. 2007, 27, 4452–4459.

40. Devor, A.; Hillman, E.; Tian, P.; Waeber, C.; Teng, I.; Ruvinskaya, L.; et al. Stimulus-induced changes in blood flow and 2-deoxyglucose uptake dissociate in ipsilateral somatosensory cortex. J. Neurosci. 2008, 28, 14347–14357.

41. Drew, P.; Shih, A.; Kleinfeld, D. Fluctuating and sensory-induced vasodynamics in rodent cortex extend arteriole capacity. Proc. Natl. Acad. Sci. 2011, 108, 8473–8478.

42. Hillman, E.; Devor, A.; Bouchard, M.; Dunn, A.; Krauss, G.; Skoch, J.; et al. Depth-resolved optical imaging and microscopy of vascular compartment dynamics during somatosensory stimulation. NeuroImage 2007, 35, 89–104.

43. Uhlirova, H.; Kıvılcım, K.; Peifang, T.; Martin, T.; Desjardins, M.; Payam, A.; et al. Cell type specificity of neurovascular coupling in cerebral cortex. eLife 2016, 5.

44. Blinder, P.; Tsai, P.; Kaufhold, J.; Knutsen, P.; Suhl, H.; Kleinfeld, D. The cortical angiome: An interconnected vascular network with noncolumnar patterns of blood flow. Nat. Neurosci. 2013, 16, 889–897.

45. Markram, H.; Muller, E.; Ramaswamy, S.; Reimann, M.; Abdellah, M.; Sanchez, C.; et al. Reconstruction and simulation of neocortical microcircuitry. Cell 2015, 163, 456–492.

46. de Kock, C.; Bruno, R.; Spors, H.; Sakmann, B. Layer- and cell-type-specific suprathreshold stimulus representation in rat primary somatosensory cortex. J. Physiol. 2007, 581, 139–154.

47. Ellsworth, M.; Ellis, C.; Popel, A.; Pittman, R. Role of microvessels in oxygen supply to tissue. News Physiol. Sci. 1994, 9, 119–123.

48. Tsai, A.; Johnson, P.; Intaglietta, M. Oxygen gradients in the microcirculation. Physiol. Rev. 2003, 83, 933–963.

49. Carlson, B.; Arciero, J.; Secomb, T.W. Theoretical model of blood flow autoregulation: Roles of myogenic, shear-dependent, and metabolic responses. Am. J. Physiol. Heart Circ. Physiol. 2008, 295, H1572–H1579.

50. Schmid, F.; Tsai, P.; Kleinfeld, D.; Jenny, P.; Weber, B. Depth-dependent flow and pressure characteristics in cortical microvascular networks. PLoS Comput. Biol. 2017, 13, e1005392.

51. Harper, S.; Bohlen, H. Microvascular adaptation in the cerebral cortex of adult spontaneously hypertensive rats. Hypertension 1984, 6, 408–419.

52. Kim, T.; Goodwill, P.; Chen, Y.; Conolly, S.; Schaffer, C.; Liepmann, D.; et al. Line-scanning particle image velocimetry: An optical approach for quantifying a wide range of blood flow speeds in live animals. PLoS ONE 2012, 7, e38590.

53. Ward, A.; Glynn, P. Properties of the reflected Ornstein–Uhlenbeck process. Queueing Syst. Theory Appl. 2003, 44, 109–123.

54. Uhlenbeck, G.E.; Ornstein, L.S. On the theory of the Brownian motion. Phys. Rev. 1930, 36, 823–841. Available online: https://link.aps.org/doi/10.1103/PhysRev.36.823 (accessed on [insert date]).

55. Brent, R.P. Algorithms for Minimization without Derivatives; Prentice-Hall: Englewood Cliffs, NJ, USA, 1973; Chapter 4.

56. Battini, S.; Cantarutti, N.; Kotsalos, C.; Carel, T. AstroVascPy. Available online: https://github.com/BlueBrain/AstroVascPy (accessed on [insert date]).

57. Dalcin, L.D.; Paz, R.R.; Kler, P.A.; Cosimo, A. Parallel distributed computing using Python. Adv. Water Resour. 2011, 34, 1124–1139. New computational methods and software tools.

58. Petersen, C.C. Sensorimotor processing in the rodent barrel cortex. Nat. Rev. Neurosci. 2019, 20, 533–546.

59. Ayaz, A.; Stäuble, A.; Hamada, M.; Wulf, M.; Saleem, A.; Helmchen, F. Layer-specific integration of locomotion and sensory information in mouse barrel cortex. Nat. Commun. 2019, 10, 2585.

60. Adesnik, H.; Naka, A. Cracking the function of layers in the sensory cortex. Neuron 2018, 100, 1028–1043.

61. Lindvere, L.; Janik, R.; Dorr, A.; Chartash, D.; Sahota, B.; Sled, J.; et al. Cerebral microvascular network geometry changes in response to functional stimulation. NeuroImage 2013, 71, 248–259.

62. Sekiguchi, Y.; Takuwa, H.; Kawaguchi, H.; Kikuchi, T.; Okada, E.; Iwao, K.; et al. Pial arteries respond earlier than penetrating arterioles to neural activation in the somatosensory cortex in awake mice exposed to chronic hypoxia: An additional mechanism to proximal integration signaling? J. Cereb. Blood Flow 2014, 34, 1761–1770.

63. Tian, P.; Teng, I.; May, L.; Kurz, R.; Lu, K.; Scadeng, M.; et al. Cortical depth-specific microvascular dilation underlies laminar differences in blood oxygenation level-dependent functional MRI signal. Proc. Natl. Acad. Sci. USA 2010, 107, 15246–15251.

64. Bélanger, M.; Allaman, I.; Magistretti, P. Brain energy metabolism: Focus on astrocyte-neuron metabolic cooperation. Cell Metab. 2011, 14, 724–738.

65. Mishra, A.; Gordon, G.; MacVicar, B.A.; Newman, E.A. Astrocyte regulation of cerebral blood flow in health and disease. Cold Spring Harb. Perspect. Biol. 2024, 16, a041354.

66. Marina, N.; Christie, I.; Korsak, A.; Doronin, M.; Brazhe, A.; Hosford, P.; et al. Astrocytes monitor cerebral perfusion and control systemic circulation to maintain brain blood flow. Nat. Commun. 2020, 11, 131.

67. Villringer, A.; Them, A.; Lindauer, U.; Einhäupl, K.; Dirnagl, U. Capillary perfusion of the rat brain cortex: An in vivo confocal microscopy study. Circ. Res. 1994, 75, 55–62.

68. Gutierrez-Jimenez, E.; Cai, C.; Mikkelsen, I.; Rasmussen, P.; Angleys, H.; Merrild, M.; et al. Effect of electrical forepaw stimulation on capillary transit-time heterogeneity (CTH). J. Cereb. Blood Flow Metab. 2016, 36, 2072–2086.

69. Shih, A.; Blinder, P.; Tsai, P.; Friedman, B.; Stanley, G.; Lyden, P.; et al. The smallest stroke: Occlusion of one penetrating vessel leads to infarction and a cognitive deficit. Nat. Neurosci. 2013, 16, 55–63.

70. Baskurt, O.; Farley, R.; Meiselman, H. Erythrocyte aggregation tendency and cellular properties in horse, human, and rat: A comparative study. Am. J. Physiol. 1997, 273, H2604–H2612.

71. Pries, A.; Secomb, T.; Gaehtgens, P.; Gross, J. Blood flow in microvascular networks: Experiments and simulation. Circ. Res. 1990, 67, 826–834.

72. Pries, A.; Secomb, T.; Gaehtgens, P.; Gross, J. Structure and hemodynamics of microvascular networks: heterogeneity and correlations. Am J Physiol. 1990, 269, H1713–22.

73. Filosa, J.; Bonev, A.; Straub, S.; Meredith, A.L.; Knot, H.J.; Nelson, M.T. Local potassium signaling couples neuronal activity to vasodilation in the brain. Nat. Neurosci. 2006, 9, 1397–1403.

74. Iadecola, C.; Nedergaard, M. Glial regulation of the cerebral microvasculature. Nat. Neurosci. 2007, 10, 1369–1376.

75. Magistretti, P.; Allaman, I. Lactate in the brain: From metabolic end-product to signalling molecule. Nat. Rev. Neurosci. 2018, 19, 235–249.

76. Gonzales, AL.; Klug, NR.; Moshkforoush, A.; J.C. Lee, JC.; Lee, FK.; Shui, B.; Tsoukias, NM.; Kotlikoff, MI.; Hill-Eubanks, D.; Nelson, MT. Proc. Natl. Acad. Sci. 2020, 117, 27022–27033.

77. Sancho, M.; Klug, NR.; Harraz, OF.; Hill-Eubanks, D.; Nelson, MT. Distinct potassium channel types in brain capillary pericytes. Biophysical Journal. 2024, 123, 2110–2121,.

78. Kleinfeld, D.; Blinder, P.; Drew, P.J.; Driscoll, J.D.; Muller, A.; Tsai, P.S.; Shih, A.Y. A Guide to Delineate the Logic of Neurovascular Signaling in the Brain. Frontiers in Neuroenergetics. 2011, 3, 1662–6427.

79. Longden T.A.; Dabertrand, F.; Koide, M.; Gonzales, AL.; Tykocki, NR.; Brayden, JE.; Hill-Eubanks, D.; Nelson, MT. Capillary K+-sensing initiates retrograde hyperpolarization to increase local cerebral blood flow. Nat Neurosci. 2017, 20, 717–726,.

80. Hartmann, D.A.; Berthiaume, A.A.; Grant, R.I.; Harrill, S.A.; Koski, T.; Tieu, T.; McDowell, K.P.; Faino, A.V.; Kelly, A.L.; Shih, A.Y.; Nelson, MT. Brain capillary pericytes exert a substantial but slow influence on blood flow. Nat Neurosci. 2021, 24, 633–645,.

81. Hill, R.A.; Tong, L.; Yuan, P.; Murikinati, S.; Gupta, S.; Grutzendler, J. Regional Blood Flow in the Normal and Ischemic Brain Is Controlled by Arteriolar Smooth Muscle Cell Contractility and Not by Capillary Pericytes. Neuron. 2015, 87, 95–110,.

82. Cauli, B.; Tong, X.K,; Rancillac, A.; Serluca, N.; Lambolez, B.; Rossier, J.; and Hamel, E. Cortical GABA Interneurons in Neurovascular Coupling: Relays for Subcortical Vasoactive Pathways. J Neurosci. 2004, 24, 8940–8949,.

83. Cauli, B., Hamel, E. Revisiting the role of neurons in neurovascular coupling. Front. Neuroenergetics. 2010, 24,,.

84. Neubauer-Geryk, J.; Hoffmann, M.; Wielicka, M.; Piec, K.; Kozera, G.; Brzeziński, M.;, Bieniaszewski, L. Current methods for the assessment of skin microcirculation: Part 1. Postepy Dermatol Alergol. 2019, 24, 247–254,.

85. Neubauer-Geryk, J.; Hoffmann, M.; Wielicka, M.; Piec, K.; Kozera, G.; Bieniaszewski, L. Current methods for the assessment of skin microcirculation: Part 2. Postepy Dermatol Alergol. 2019, 36, 377–381,.

86. Broggini, T.; Duckworth, J.; Ji, X.; Liu, R.; Xia, X.; Machler, P.; Shaked, I.; Munting, LP.; Iyengar, S.; Kotlikoff, M.;7 van Veluw, S.J.; Vergassola, M.; Gal Mishne, G.; David Kleinfeld, D. Long-wavelength traveling waves of vasomotion modulate the perfusion of cortex. Neuron. 2024, 112, 2349–2367,.

87. Attwell, D.; Laughlin, S. An energy budget for signaling in the grey matter of the brain. J. Cereb. Blood Flow Metab. 2001, 21, 1133–1145.

88. Hossmann, K. Viability thresholds and the penumbra of focal ischemia. Ann. Neurol. 1994, 36, 557–565.

89. Schulte, M.; Wood, J.; Hudetz, A. Cortical electrical stimulation alters erythrocyte perfusion pattern in the cerebral capillary network of the rat. Brain Res. 2003, 963, 81–92.

90. Stefanovic, B.; Hutchinson, E.; Yakovleva, V.; Schram, V.; Russell, J.; Belluscio, L.; et al. Functional reactivity of cerebral capillaries. J. Cereb. Blood Flow Metab. 2008, 28, 961–972.

91. Kleinfeld, D.; Mitra, P.; Helmchen, F.; Denk, W. Fluctuations and stimulus-induced changes in blood flow observed in individual capillaries in layers 2 through 4 of rat neocortex. Proc. Natl. Acad. Sci. USA 1998, 95, 15741–15746.

92. Hutchinson, E.; Stefanovic, B.; Koretsky, A.; Silva, A. Spatial flow-volume dissociation of the cerebral microcirculatory response to mild hypercapnia. NeuroImage 2006, 32, 520–530.

93. Unekawa, M.; Tomita, M.; Tomita, Y.; Toriumi, H.; Miyaki, K.; Suzuki, N. RBC velocities in single capillaries of mouse and rat brains are the same, despite 10-fold difference in body size. Brain Res. 2010, 1320, 69–73.

94. Schmid, F.; Barrett, M.; Jenny, P.; Weber, B. Vascular density and distribution in neocortex. NeuroImage 2019, 197, 792–805.

95. Epp, R.; Schmid, F.; Weber, B.; Jenny, P. Predicting vessel diameter changes to up-regulate biphasic blood flow during activation in realistic microvascular networks. Front. Physiol. 2020, 16, e1005392.

96. Vaucher, E.; Hamel, E. Cholinergic basal forebrain neurons project to cortical microvessels in the rat: Electron microscopic study with anterogradely transported Phaseolus vulgaris leucoagglutinin and choline acetyltransferase immunocytochemistry. J. Neurosci. 1995, 15, 7427–7441.

97. Hartung, G.; Vesel, C.; Morley, R.; Alaraj, A.; Sled, J.; Kleinfeld, D.; et al. Simulations of blood as a suspension predicts a depth-dependent hematocrit in the circulation throughout the cerebral cortex. PLoS Comput. Biol. 2018, 14, e1006549.

98. Gould, I.; Linninger, A. Hematocrit distribution and tissue oxygenation in large microcirculatory networks. Microcirculation 2015, 22, 1–18.

99. Gould, I.; Tsai, P.; Kleinfeld, D.; Linninger, A. The capillary bed offers the largest hemodynamic resistance to the cortical blood supply. J. Cereb. Blood Flow Metab. 2017, 37, 52–68.

100. Ozawa, K.; Nagao, M.; Konno, A.; Iwai, Y.; Vittani, M.; Kusk, P.; et al. Astrocytic GPCR-induced Ca2+ signaling is not causally related to local cerebral blood flow changes. Int. J. Mol. Sci. 2023, 24, 13590.

101. Clavreul, S.; Dumas, L.; Loulier, K. Astrocyte development in the cerebral cortex: Complexity of their origin, genesis, and maturation. Front. Neurosci. 2022, 16.

102. Tan, C.; Bindu, D.; Hardin, E.; Sakers, K.; Baumert, R.; Ramirez, J.; et al. δ-Catenin controls astrocyte morphogenesis via layer-specific astrocyte-neuron cadherin interactions. J. Cell Biol. 2023, 222, 11.

103. Ji, X.; Ferreira, T.; Friedman, B.; Liu, R.; Liechty, H.; Bas, E.; et al. Brain microvasculature has a common topology with local differences in geometry that match metabolic load. Neuron 2021, 109, 1168–1187.

104. Xiong, B.; Li, A.; Lou, Y.; Chen, S.; Long, B.; Peng, J.; et al. Precise cerebral vascular atlas in stereotaxic coordinates of whole mouse brain. Front. Neuroanat. 2017, 11, 128.

105. Paulson, O.B.; Strandgaard, S.; Edvinsson, L. Cerebral autoregulation. Cerebrovasc. Brain Metab. Rev. 1990, 2, 161–192.

106. Budohoski, K.P.; Czosnyka, M.; Kirkpatrick, P.J.; Smielewski, P.; Steiner, L.A.; Pickard, J.D. Clinical relevance of cerebral autoregulation following subarachnoid haemorrhage. Nat. Rev. Neurol. 2013, 9, 152–163.

