## Supplementary figures and images for "Modeling of blood flow dynamics in the rat somatosensory cortex"

### Supplemental Figure 1

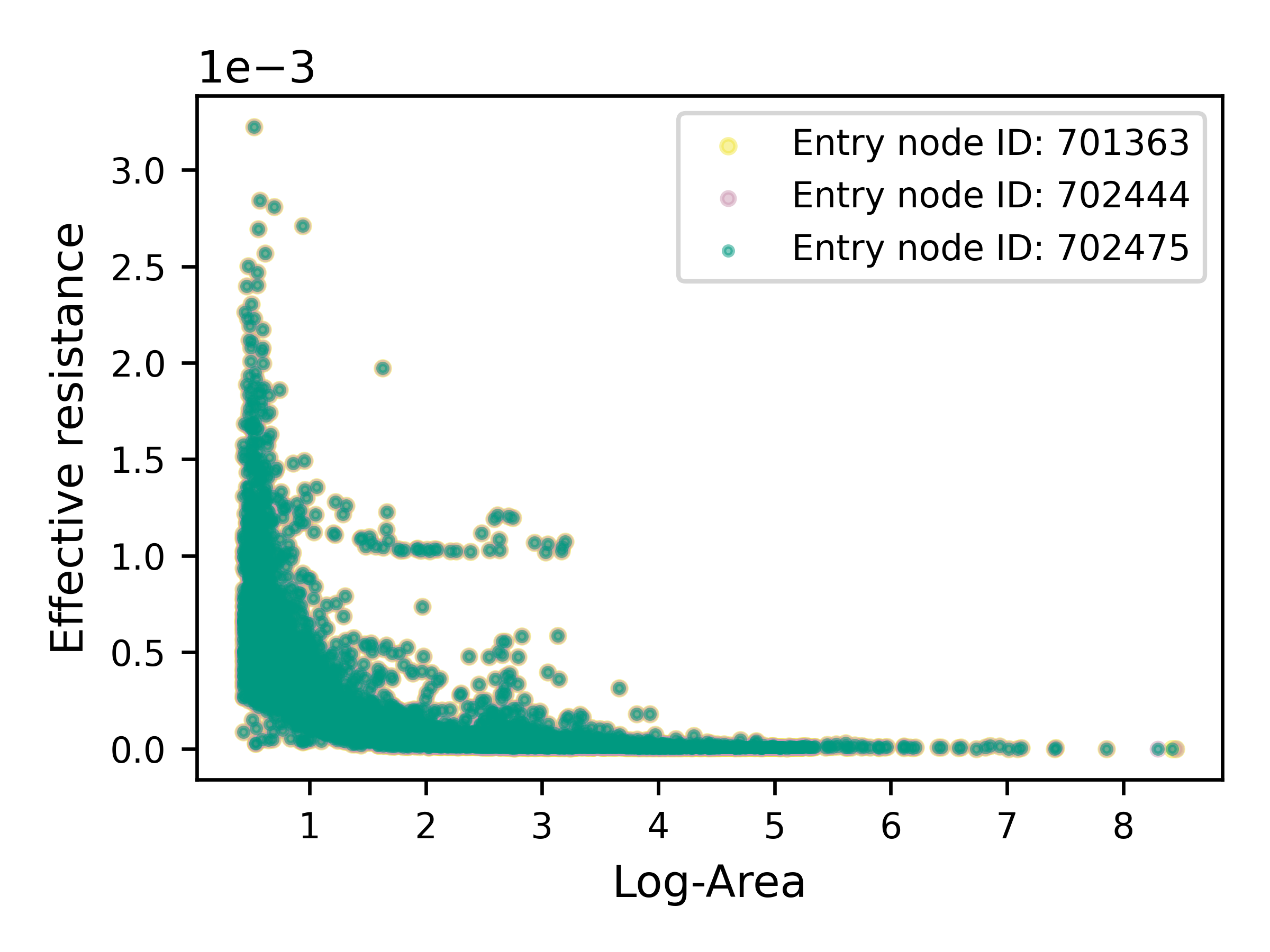

### Supplemental Figure 2

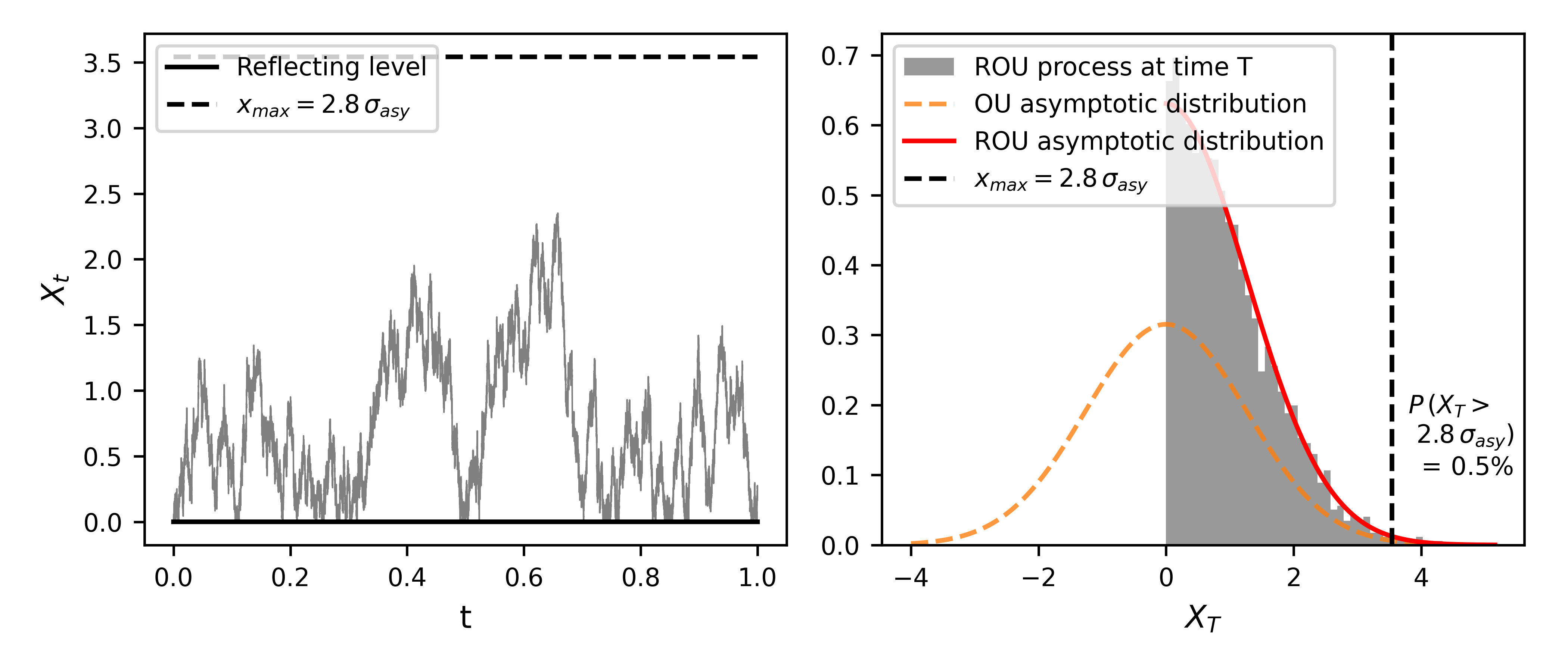

### Supplemental Figure 3

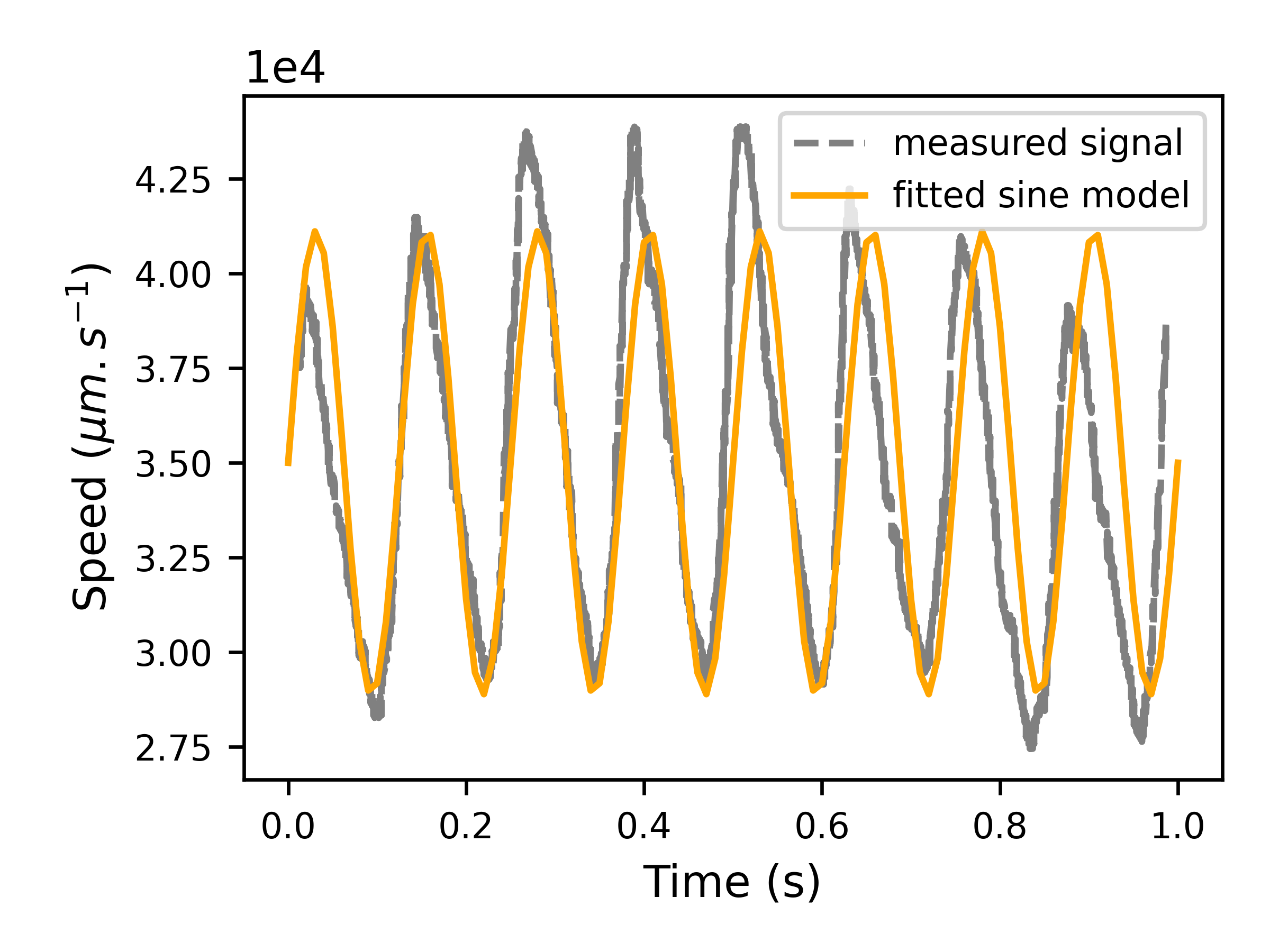

### Supplemental Figure 4

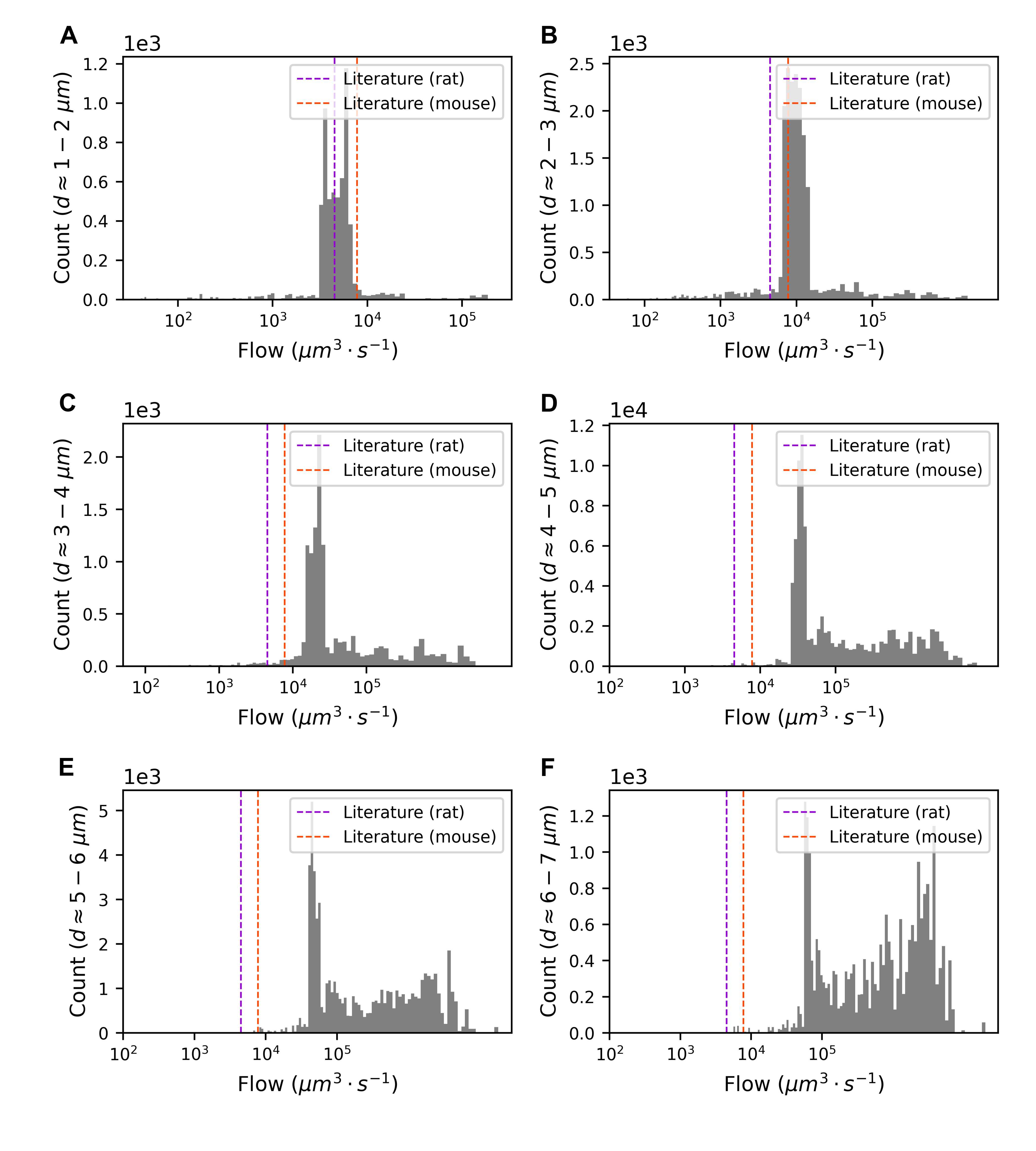

### Supplemental Figure 5

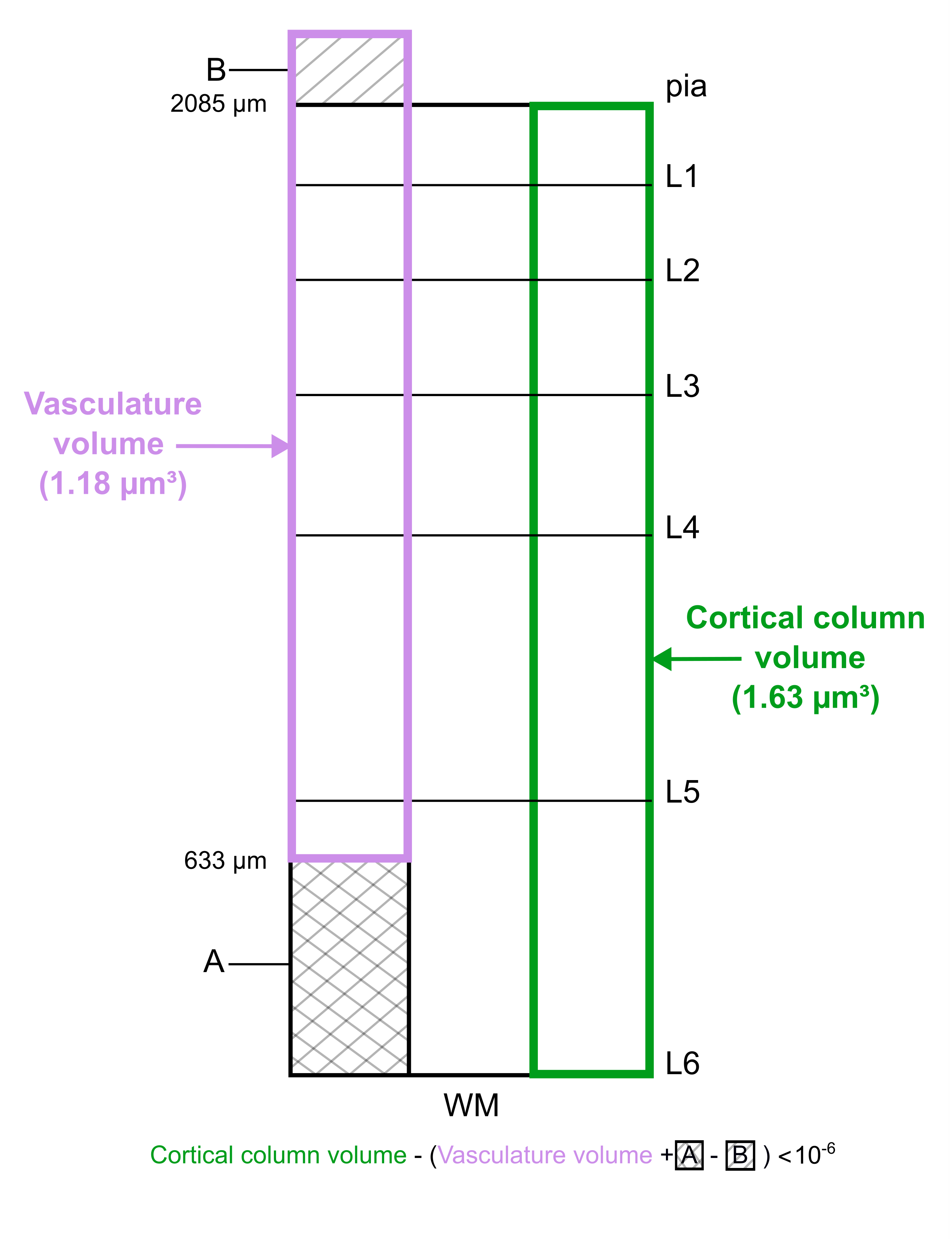

### Supplemental Figure 6

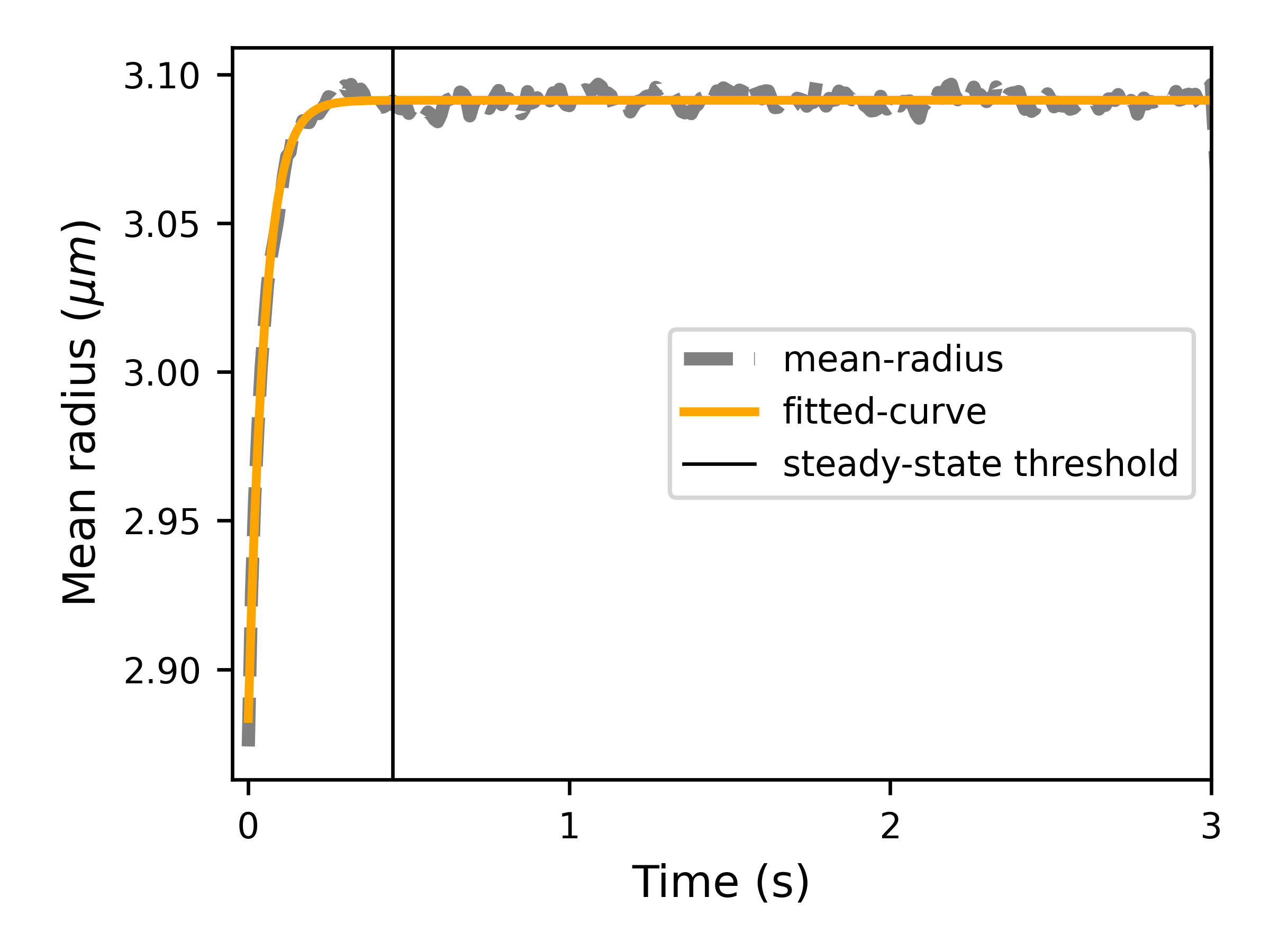

### Supplemental Table 1

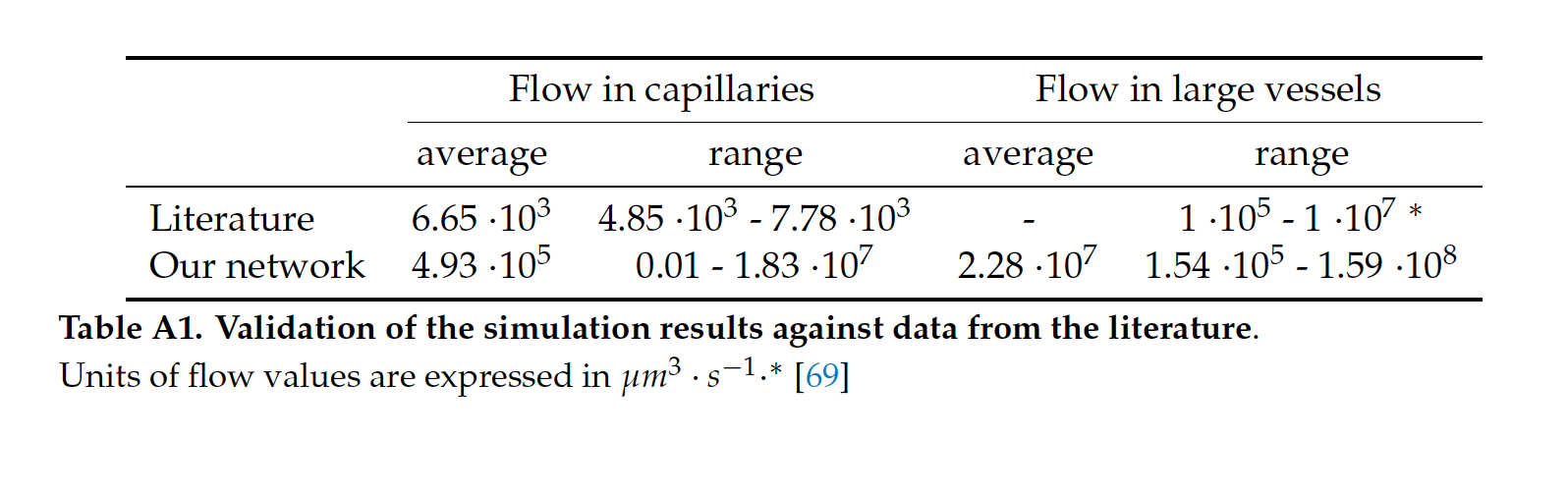

### Supplemental Table 2

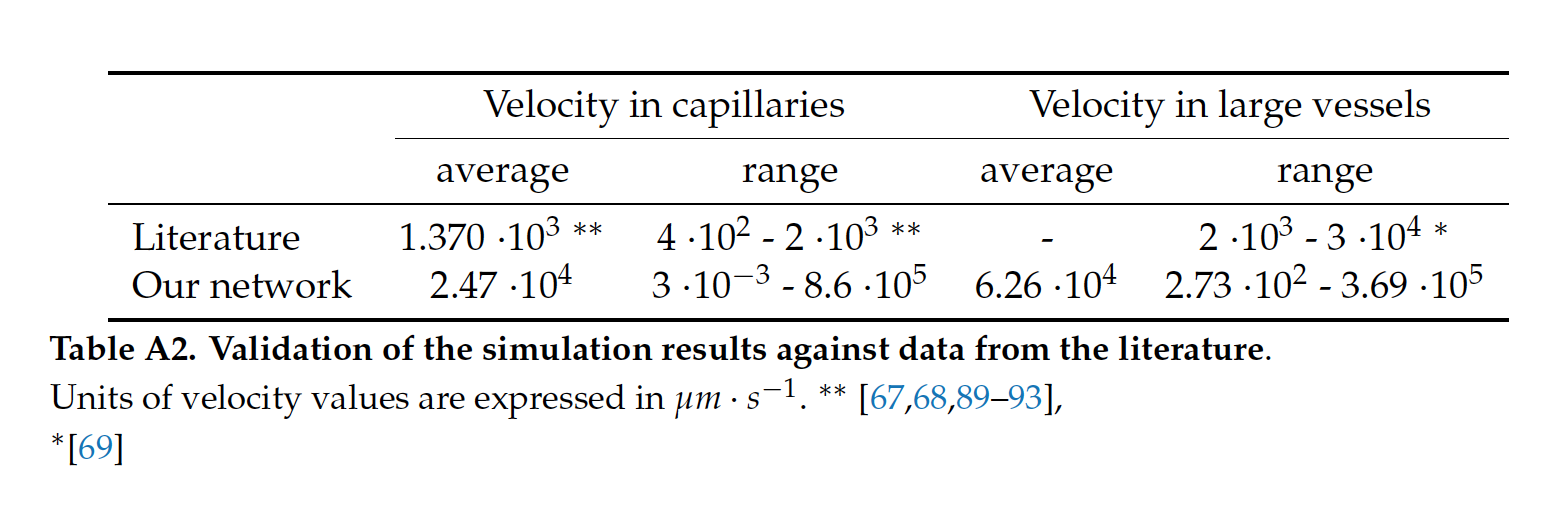

### Supplemental Table 3

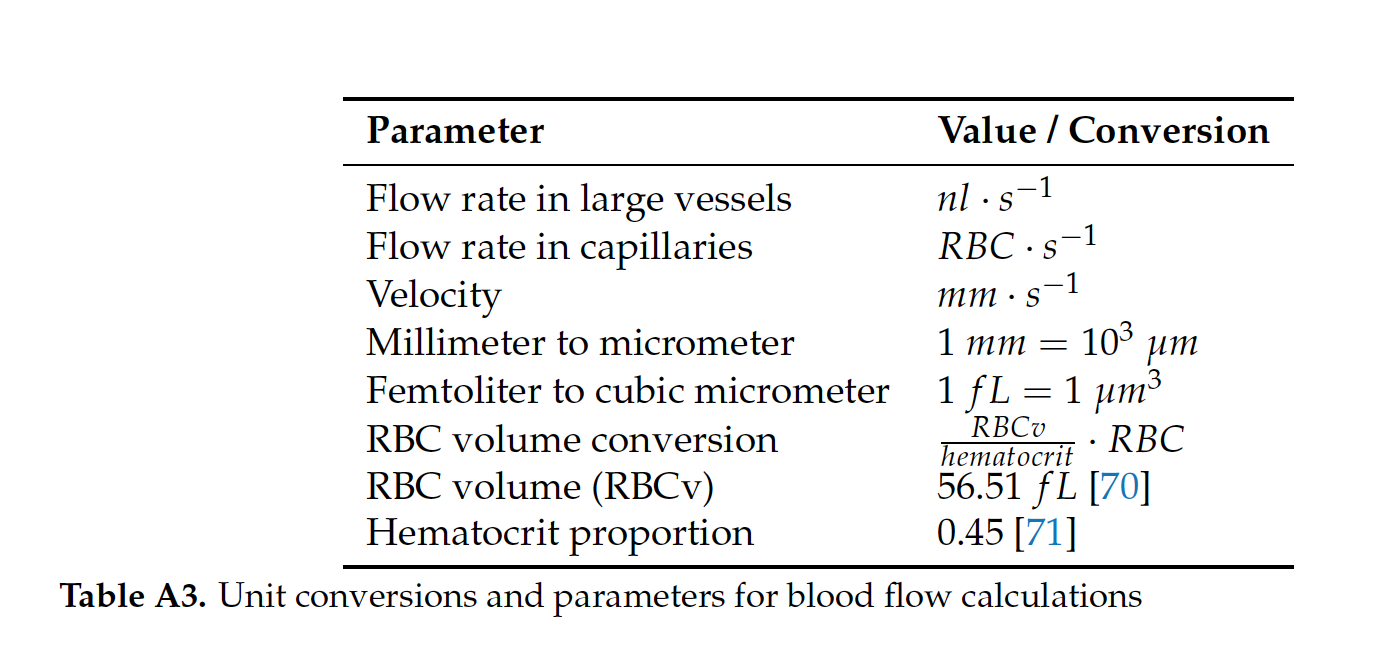
